# Public RNA-seq data are not representative of global human diversity

**DOI:** 10.1101/2024.10.11.617967

**Authors:** Irene Gallego Romero, Grace Rodenberg, Audrey M. Arner, Lani Li, Isobel J. Beasley, Ryan Rossow, Nicholas Ryan, Selina Wang, Amanda J. Lea

## Abstract

The field of human genetics has reached a consensus that it is important to work with diverse and globally representative participant groups. This diverse sampling is required to build a robust understanding of the genomic basis of complex traits and diseases as well as human evolution, and to ensure that all people benefit from downstream scientific discoveries. While previous work has characterized compositional biases and disparities for public genome-wide association (GWAS), microbiome, and epigenomic studies, we currently lack a comprehensive understanding of the degree of bias for transcriptomic studies. To address this gap, we analyzed the metadata for RNA-seq studies from two public databases—the Sequence Read Archive (SRA), representing 795,071 samples from 21,209 studies, and the Database of Genotypes and Phenotypes (dbGaP), representing 167,389 samples from 649 studies. We also randomly selected 620 studies from SRA for detailed, manual evaluation. We found that 3% of samples in SRA and 21% of individuals described in the literature had population descriptors (race, ethnicity, or ancestry); 28% of samples in dbGaP had paired genotype data that was used to empirically infer ancestry. In SRA, dbGaP, and the literature, race, ethnicity, and ancestry terms were frequently conflated and difficult to disambiguate. After standardizing population descriptors, we observed many clear biases: for example, among samples in SRA that were coded using US Census terms, 69.0% came from white donors, corresponding to an 1.2x overrepresentation of this group relative to the US population. Among samples in SRA coded using continental ancestry labels, 55.6% came from European ancestry donors—an 4.1x overrepresentation of this group relative to the global population. These biases were generally similar across datasets (SRA, dbGaP, literature review), and were comparable to previous reports for other ‘omics data types. However, we note that, relative to other ‘omics data subsets like GWAS, there is considerably less information, of arguably worse quality, about who is participating in RNA-seq studies. Together, these results demonstrate a critical need to improve our thoughtfulness, consistency, and effort around reporting population descriptors in RNA-seq studies, and to more generally strive for greater diversity in this important data type.

## INTRODUCTION

Research that includes diverse populations is essential for building a better understanding of human disease, biology, and evolution. It is also essential for combating health disparities and ensuring that all people benefit from ongoing scientific discoveries [1,2]. A growing body of work has highlighted the lack of environmental, geographic, racial, ethnic, and genetic diversity in human biomedical and especially human genomic studies. For example, an estimated 80% and 74% of participants in biomedical and microbiome studies, respectively, are from high-income countries, while an estimated 95% and 87% of participants in genome-wide association (GWAS) and epigenomic studies are of European ancestry [3–6]. These biases stem from many complex factors, including global funding patterns, legacies of culture-specific racism and exclusion toward certain identities, as well as a historical lack of reciprocity, trust, and transparency between underrepresented groups and biomedical researchers [7,8].

Overcoming these issues will require multi-faceted approaches, but as a starting point we also need a comprehensive description of the problem. While previous work has generated these descriptions for certain ‘omics data and analysis types (e.g., microbiome data [3] and GWAS [6,9]), less work has focused on functional genomic data. Previous attempts in this area have analyzed the composition of key consortia datasets (e.g., TCGA [10] or ENCODE [5]) or certain analysis types (e.g., eQTL or EWAS [10]). However, we still lack a comprehensive understanding of reporting and representation in gene expression studies more broadly.

The transcriptome is a complex cellular phenotype that has served as a model for understanding the genetic and environmental basis of complex traits and diseases, aging and developmental processes, and human ecology and evolution [11]. Over the past decade, RNA-seq approaches in particular (e.g., mRNA-seq, scRNA-seq, as well as other technologies) have become deeply integrated into many disciplines. For example, a large body of work has paired genotype and RNA-seq data to uncover the genetic architecture of gene expression via expression quantitative trait mapping (eQTL), with an increasing appreciation that these genetic effects vary across ancestry groups due to differences in genomic patterns (e.g., allele frequency, linkage disequilibrium) as well as tissue identity, pathogen exposure, or other aspects of environment [12–16]. In addition to serving as models for genetic effects, transcriptomic studies have also been instrumental in uncovering how key aspects of our environment influence health, for example showing that social determinants like socioeconomic status, social integration, or rural-urban gradients impact transcription in circulating blood [17–19]. Thus, gene expression variation has clearly been linked to genotype as well as social, economic, and ecological factors that vary worldwide, reinforcing the need to build diverse and representative transcriptomic datasets with appropriate and well-curated information on genetics and environment.

Presently, publications and public databases typically use one or multiple population descriptor terms to describe study participants, including race and ethnicity (sociopolitical constructs), continental ancestry group (to capture a person’s broad descent or heritage), nationality, and/or geographic location (to capture a person’s broad environment, but often also used as a proxy for ancestral background) [20] (**Table S1**). These descriptors reflect imperfect concepts that categorize people into groups according to self-identified, perceived, or evaluated characteristics; in many cases, these groupings were originally meant to act as stand-ins for genetic similarity between individuals, a task for which they are ill-suited. In other words, population descriptors poorly capture the complex and continuous patterns of human genetic variation and should be used with care, in service to the specific study goals [20]. Thus, while we believe the ultimate goal of human genetics should be to operationalize our complex and continuous reality, here we describe the breakdown of public RNA-seq studies using existing racial, ethnic, continental ancestry, and geographic terms. In doing so, we aim to document and highlight biases in our understanding and representation of human transcriptomic variation.

To do so, we downloaded the metadata for 21,209 human RNA-seq studies representing 1,121,832 submitted samples from the Sequence Read Archive (SRA), as well as 649 studies representing 167,389 unique samples from the Database of Genotypes and Phenotypes (dbGaP). We analyzed the degree to which 1) population descriptor terms (race, ethnicity, ancestry, geography) are reported, 2) the breakdown of those terms, and 3) other aspects of bias such as which countries, tissues, and diseases are most well-represented. Importantly, we repeated our main analyses after performing a manual review of 620 randomly selected SRA studies to understand consistency between what is reported in the literature versus what is uploaded to public databases. Our analyses reveal widespread bias in the racial, ethnic, continental ancestry, and geographic composition of current RNA-seq studies, and more generally a lack of consistent and careful reporting that needs urgent improvement.

## RESULTS

### Population descriptors are often not included or are erroneously described

We used three main approaches to understand the reporting and composition of human RNA-seq studies, focusing on studies deposited in public repositories with sample sizes ≥10, that did not focus entirely on commercially available cell lines, and that were released between 2009 and 2023. First, we used the SRA Run Selector to download all available metadata for 21,209 human RNA-seq studies, covering 1,121,832 entries for 795,071 unique “BioSamples” that met our inclusion criteria (Methods). We note that BioSample is an ambiguous descriptor, as it can refer to different individuals, different technical replicates, or to the same individual evaluated across multiple experimental conditions. In our second approach, we downloaded metadata for the 649 RNA-seq studies (covering 167,389 samples) available in dbGaP and extracted the “ancestry” term, which is empirically derived from genotype data via the GrafPop software and draws from a set of mixed continental ancestry and racial descriptors [21] (see **Table S2**). Finally, we manually reviewed a random sample of 620 studies associated with SRA depositions, and extracted a wide range of information about population descriptors and study design (**Table S3**).

In SRA, 23,896 BioSamples (3% of all BioSamples passing filters) from 261 studies were associated with some kind of population descriptor (i.e., they had data in a column labeled as ethnicity, race, ancestry, or geography; **Table 1**). After harmonization and backcoding of descriptor data, we were able to further classify these as either US census racial or ethnic terms (14,057 samples) or geographic/continental ancestry (9,839 samples) terms (Methods, **Table S4, S5**). Category and individual term usage varied across countries, with 58.8% of samples deposited into SRA by institutions outside the US using geographic/continental ancestry terms and 87.5% of samples deposited by US institutions using racial or ethnic terms. This result is somewhat unsurprising given that race is still used for legal and official purposes in the US, for example the census and in health records. However, most other countries do not recognize racial categories and have instead moved to enumerate individuals by self-reported ethnic or ancestry descriptors. In dbGaP, 28% of samples (47,022/167,389 passing filters) included empirically-derived descriptor terms (the only available/allowable option for dbGaP).

**Table 1.** Datasets included and reporting of population descriptors. Numbers in parentheses represent proportions, with numbers from the previous column as the denominator.

|  | Studies included | Analysis units | Units with any population descriptor | Units with geographic/continental ancestry | Units with US Census term |
| --- | --- | --- | --- | --- | --- |
| SRA | 21,209 | 795,071 BioSamples | 23,896 (0.03) | 9,839 (0.41) | 14,057 (0.59) |
| dbGaP | 649 | 167,389 individuals | 47,022 (0.28) | 47,022 (1) | 0 |
| Literature Review | 312 | 16,801 individuals | 3,376 (0.20) | 2,126 (0.63) | 1,250 (0.37) |

We randomly selected 620 SRA accessions and manually evaluated each associated publication in detail for reporting of any population descriptors. After excluding those that did not have an available publication or only used commercial cell lines, we were left with 312 studies (n=16,801 individuals); we note that 6% of the retained 312 studies focused on LCL, fibroblast, or iPSC lines instead of primary samples. In the literature, we observed better reporting relative to what is stored in SRA, with 20% of studies (61/312) including information about participants’ race, ethnicity, or ancestry. Consistent with our observations from SRA, the majority of authors reporting US census terms were at institutions based in the United States (100%; 22/22). Given the general push to move away from categorical to continuous and data-driven representations of ancestry, we also recorded how often genotype data were available. We found that some type of genome-wide genotype data were available for 32% (101/312) of manually reviewed studies.

### RNA-seq studies are biased toward high income countries

We next focused on understanding the geographic distribution of institutions performing RNA-seq studies as well as the locations where samples were collected (although sample collection information was only available for the manually reviewed studies). Using the “Center Name” category from SRA, which typically reflects the lead institution responsible for depositing the data, we found that 62.1% of samples with population descriptors were deposited by institutions in the US, and 97.6% of samples were deposited from a High Income Country (HIC), as defined by the World Bank (**Figure 1A, 1B; Table S6**). This figure rose to 100% of samples and studies when considering samples in dbGaP, all of which were deposited by institutions in the US. We note this strong bias for dbGap is likely driven by the fact that, unlike SRA, it is not connected to external databases hosted outside of the US (e.g., DNA Data Bank of Japan or the European Bioinformatics Institute).

**Figure 1:**
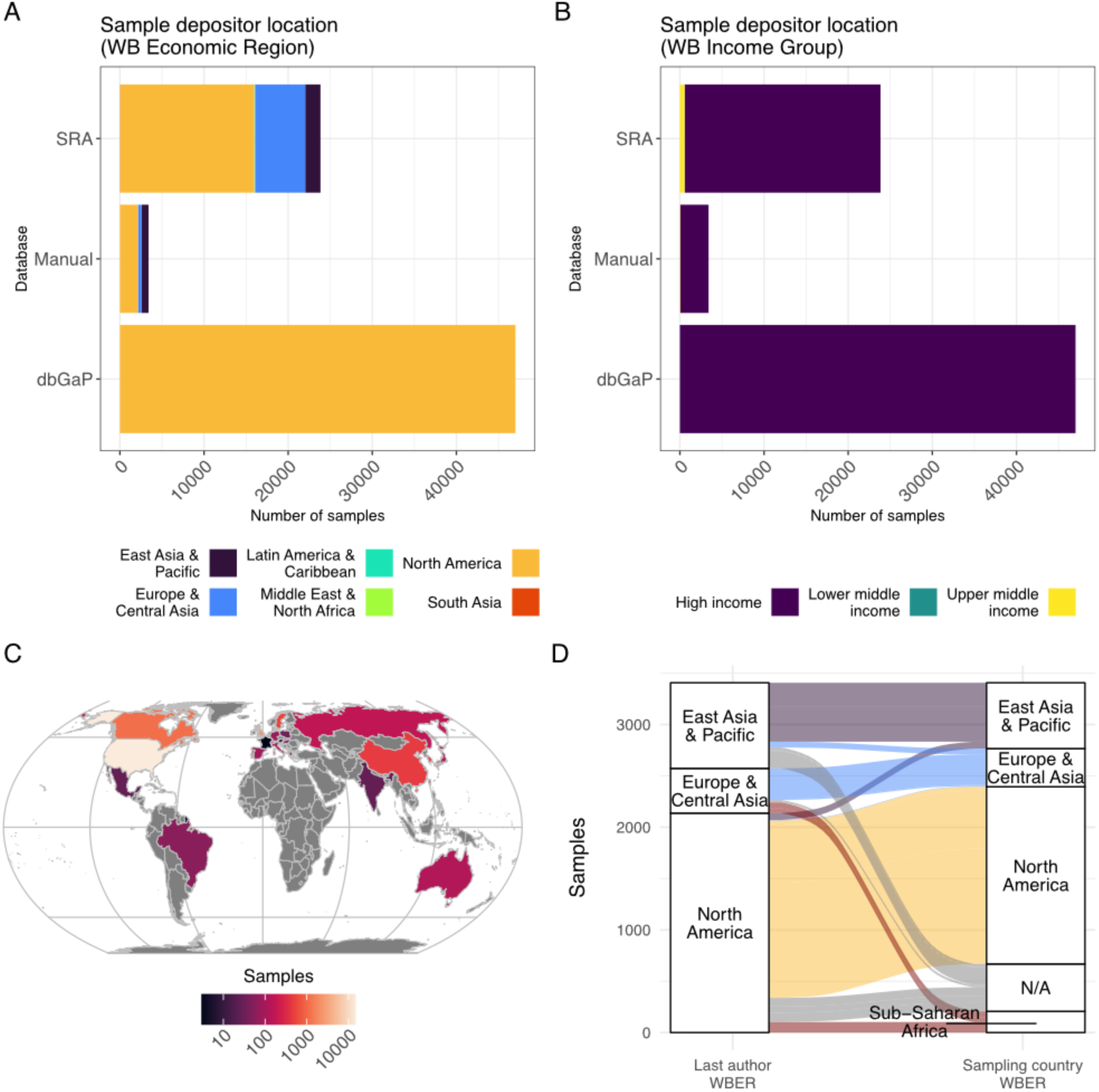
Geographic and economic features of institutions depositing transcriptomic datasets to public databases. A) Number of samples deposited to SRA, dbGaP, and in our manual review dataset, grouped by World Bank (WB) Economic Region. B) Number of samples deposited to SRA, dbGaP, and in our manual review, grouped by World Bank Income Group. C) Countries depositing transcriptomic data with population descriptors to the SRA. Each country is shaded by the number of samples it deposited, with darker colors indicating fewer samples than lighter colors. Countries represented by 0 samples are in gray. D) Location of the last author and country where samples were collected for 3,704 samples in the manual review, grouped by World Bank Economic Region (WBER). N/A represents studies where the source country for the RNA-seq study participants was unreported.

We arrived at similar conclusions about biases in where lead institutions are located via our manual review of publications: using the last author’s institutional affiliation, we found that 58% (182/312) of studies were led by institutions in the US, while 94% (292/312) of studies were led by an institution in a high income country. Only 65% (202/312) of manually reviewed studies reported information about samples’ geographic origin in publications. However, when this information was included, we found that 57% (1,675/2917) of participants were from the US. Only 12% (5/43) of studies that included geographic information focused exclusively on individuals sampled outside of HICs.

To better examine patterns of under- and overrepresentation in samples with population descriptors by country, we compared the proportion of samples deposited by institutions in a given country to the country’s relative population size (using United Nations estimates from 2020, following [3]; **Table S7, Figure S1-2**). Not surprisingly, HICs are dramatically overrepresented relative to their population sizes: for example, the US is home to approximately 4.3% of the global population, and is thus overrepresented by a factor of 14.5 in SRA and our manually curated dataset. We also found that certain regions of the world are severely underrepresented; for example, no samples were deposited by research institutions based anywhere in Africa, and only 0.3% of samples were deposited by institutions in Latin America or the Caribbean (**Figure 1C**). This does not, however, mean that individuals from these regions are completely absent from transcriptomic datasets. For example, in the manual review, which is the only dataset for which we can separate the country where samples were collected from that where the last author was based, we found that 6.3% of samples were collected in sub-Saharan Africa (**Figure 1D**).

### RNA-seq studies are biased toward White and European ancestry individuals

We then asked about the specific population descriptors used by studies to describe samples. European individuals make up the majority of samples with ancestral/geographic descriptors amongst all three datasets (**Figure 2A-C**). In the SRA dataset they account for 56% of samples (5,469/9,839), whereas in dbGaP, 63% of samples (29,607/47,022) were labeled as ‘European’ by GrafPop. Amongst the manually reviewed studies, individuals with European ancestry comprised 51% (1,076/2,126) of individuals with geographic/continental ancestry descriptors. Thus, across all three datasets, individuals of European ancestry are substantially overrepresented relative to their share of the global population—a similar trend to that observed in GWAS studies—although the excess in transcriptomic datasets is less pronounced (3.6-fold in SRA and 3.4-fold in the manually reviewed studies, **Figure S3**) [6].

**Figure 2:**
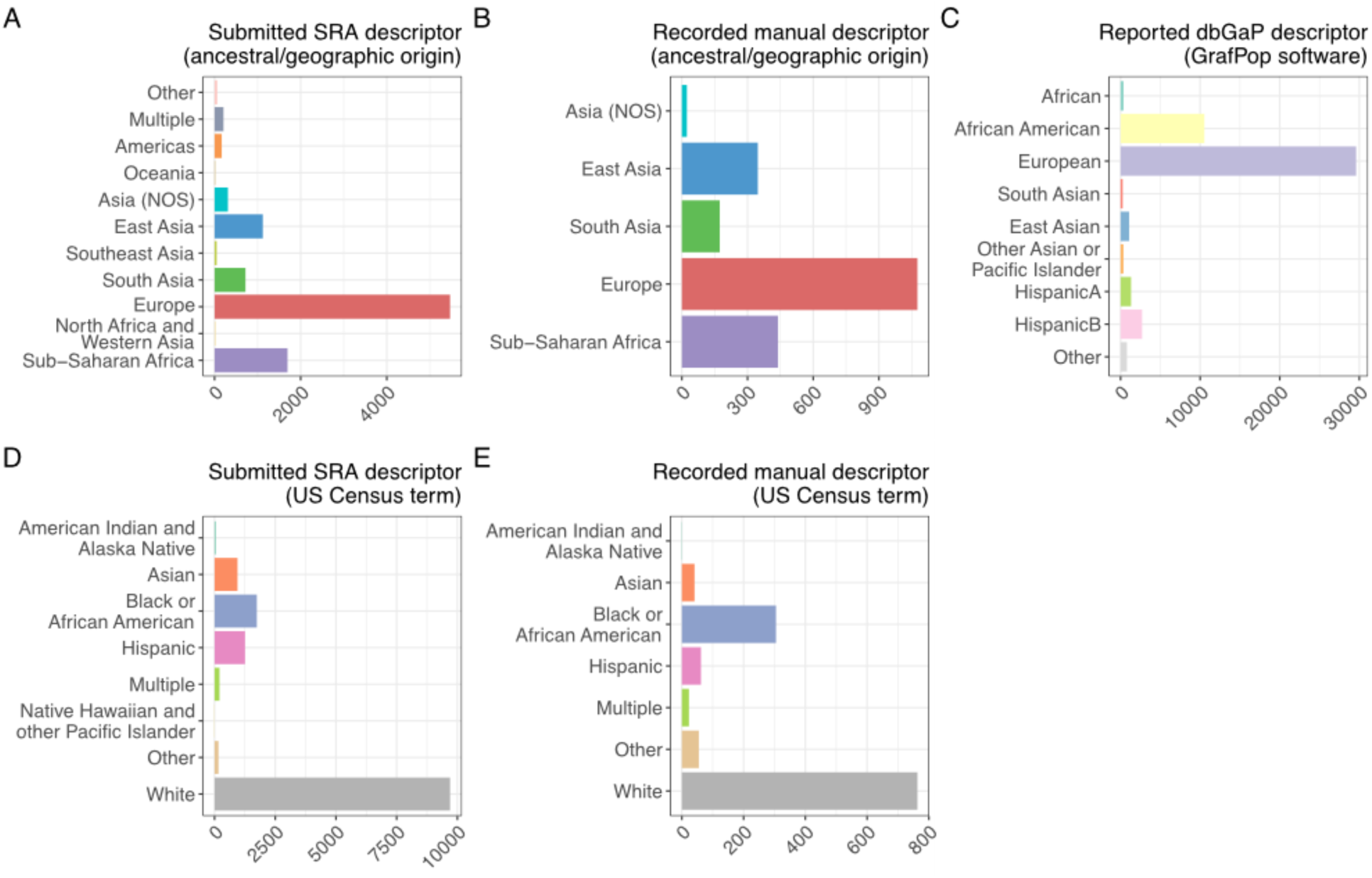
Population descriptor usage across datasets. A, B) Geographic/ancestral descriptor usage in SRA and manually reviewed studies. NOS = Not Otherwise Specified. C) GrafPop descriptors used in dbGaP studies. D, E) US Census term usage in SRA and manually reviewed studies. Asia (NOS): Asia, not otherwise specified. In all panels, the x-axis represents the number of samples.

When focusing on samples labeled with US Census terms, we observed that the predominant label was white individuals (**Figure 2D, 2E**). They accounted for 69.0% (9,700/14,055) and 61% (763/1,250) of individuals in the SRA and the manually reviewed studies respectively. Although these percentages are higher than the number of individuals who declared themselves white in the 2020 census (57% of the population, **Table S4**), amongst both the SRA and the manual datasets, the most overrepresented group of individuals are those reported as “Other” (2.26-fold in SRA, 8.63-fold in the manual data; **Figure S4**). Hispanic and multiracial individuals are consistently underrepresented in both datasets, while only the SRA includes individuals described as either American Indian and Alaska Native (43 samples, 0.3%) or Native Hawaiian and Other Pacific Islander (6 samples, 0.04%).

In both the SRA and dbGaP, we noted that European ancestry samples, and samples from white individuals, were included in more studies and had larger sample sizes per study (**Figure S5**). For instance, of the 135 studies that used geographic/ancestry descriptors in the SRA, only five included samples from Oceania. Of these, the largest study included 3 samples, and the total number of samples across studies is only 11. In contrast, 87 studies in the SRA included at least one sample of European ancestry (mean = 62.9 samples per study, max = 753 samples). We observe similar trends in the 139 SRA studies that use US Census descriptors, with the least represented group, Native Hawaiian and Other Pacific Islander, being included in only 3 studies, while white individuals were included in 106 studies. We also stratified samples by where in the world they were deposited into the SRA from, again using World Bank Economic Regions, and found that studies deposited from institutions in East Asia and the Pacific were focused primarily on Asian individuals (**Figure S6**), likely reflecting local research priorities and health concerns. Finally, we note that in all three datasets, race, ethnicity, and ancestry were often conflated, with considerable ambiguity and inconsistency in term usage both within and across studies. This reflects global differences in the usage of population descriptors (for example, “white” and “black” are racial descriptors in the US, but ethnicity descriptors in the UK) and, likely, confusion amongst researchers as to the differences between descriptor categories. Indeed, even in the manually-evaluated publications, authors conflated ancestry, ethnicity, and race. Given these ambiguities, in our backcoding we deferred to author-supplied metadata categories for grouping sample terms even when terms appeared to be incorrectly categorized (Methods).

### Other aspects of bias and composition in public RNA-seq studies

We were able to retrieve tissue of origin information for all SRA samples with population descriptor terms, which allowed us to investigate other features of study design. A total of 46 tissues are present in the dataset, although only 17 of these exceeded 100 observations combined amongst both categories of population descriptors (**Figure 3**, **Figure S7, Table S8**). Blood is the most abundant tissue amongst samples described with either geographic/ancestral terms (5,076 samples, 51.6%) or US Census terms (8,955 samples, 63.7%). Perhaps because it is easier to collect than other types of samples, we find that blood samples consistently exhibit the greatest diversity amongst all tissue types, both in terms of the institutions they are deposited from and the individuals represented (**Figures S8-9**). For example, there is at least one blood sample associated with all of the US Census descriptors, and with all geographic/ancestral descriptors except for Oceania. The diversity represented within blood samples differs significantly from that of the overall dataset (chi-square p<10^-16^ for either descriptor type, namely US Census or geographic/ancestral terms).

**Figure 3:**
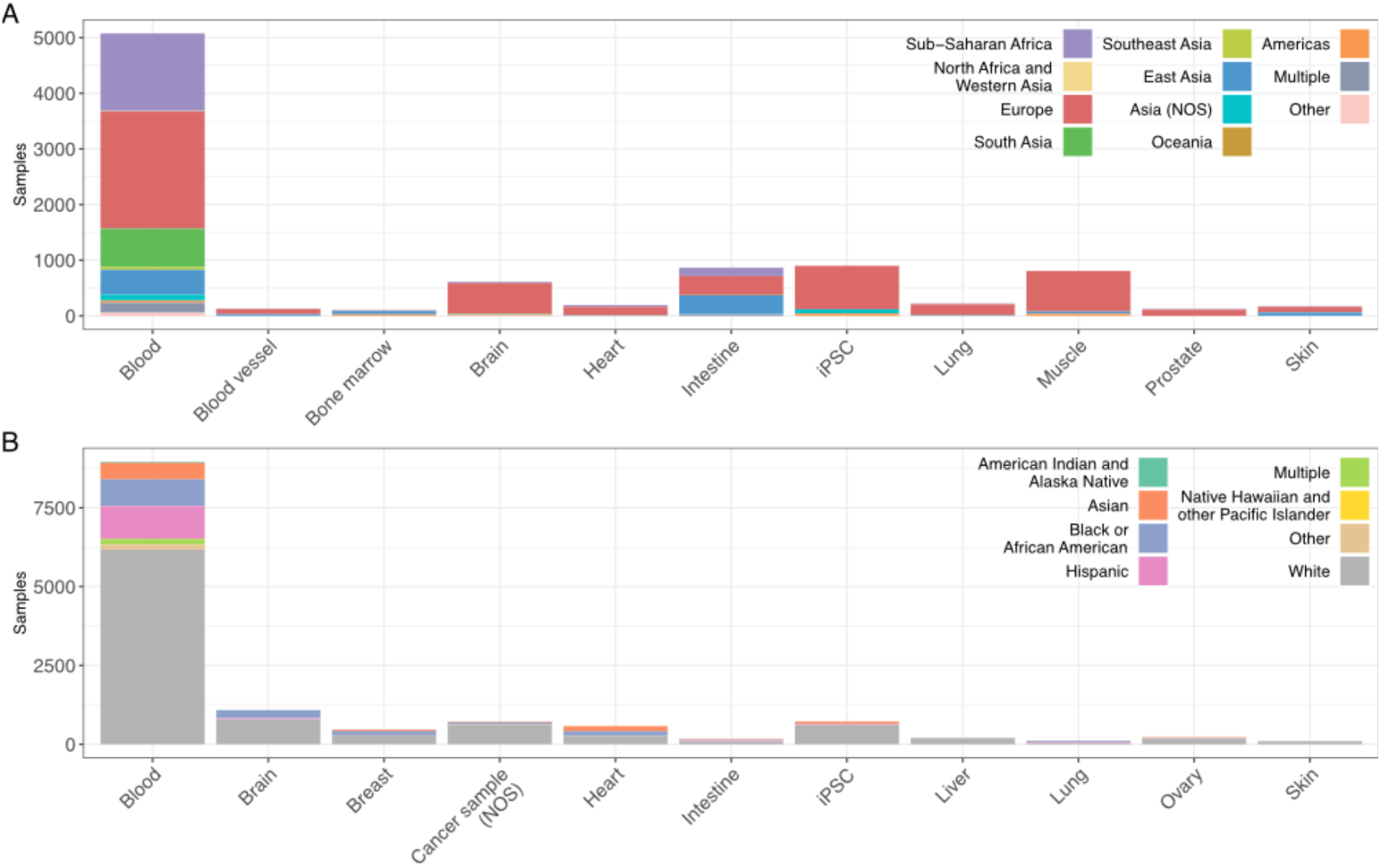
Number of individuals by population descriptor term and tissue in SRA. Number of samples per tissue (x-axis) broken down by A) Geographic/ancestral descriptors and B) US Census terms. Both panels focus on tissues with over 100 entries in SRA. A version of this plot showing all tissues is available as **Figure S7**.

When analyzing tissue diversity within ancestry or racial groups, tissue representation is greatest amongst individuals of European ancestry (23 tissues, 82.1% of all tissues with at least one sample with a geographic/ancestral descriptor) and white individuals (36 tissues, 97.3% of all tissues with at least one sample with a US Census descriptor). Individuals from sub-Saharan Africa and Black or African American individuals were the second-best represented class (16 and 25 tissues respectively), but multiple groups (Oceania, Southeast Asia, and Native Hawaiian and other Pacific Islanders) were represented by only one tissue type. This observation highlights a novel and underappreciated axis of disparity in publicly available transcriptomic datasets.

Using information provided in SRA, we were able to identify the disease focus of 6,161 samples with population descriptors (3,050 and 3,111 samples with geographic and census descriptors; **Figure 4, Figure S10, Table S9**). This number includes samples described as “healthy controls”, which is the most common label amongst samples associated with geographic descriptors (n = 1,134) and the second-most common amongst those labeled with US Census terms (n = 930). This data enabled us to explore the relationships between population descriptor usage, SRA depositing institution, and tissue and disease focus (**Figure S11-12**). For example, regardless of descriptor usage, 907 cancer samples (59.2% of all cancer samples) were deposited to the SRA by institutions in North America, followed by 380 (24.8%) and 208 (13.6%) samples from institutions located in East Asia and the Pacific, and in Europe and Central Asia, respectively. This bias likely reflects cancer’s status as a research focus and healthcare priority worldwide. In contrast, of the 983 samples with an autoimmune disease focus, 852 (86.7%) were deposited by North American institutions, potentially reflecting the fact that countries with fewer healthcare and research resources do not prioritize this set of disorders.

**Figure 4:**
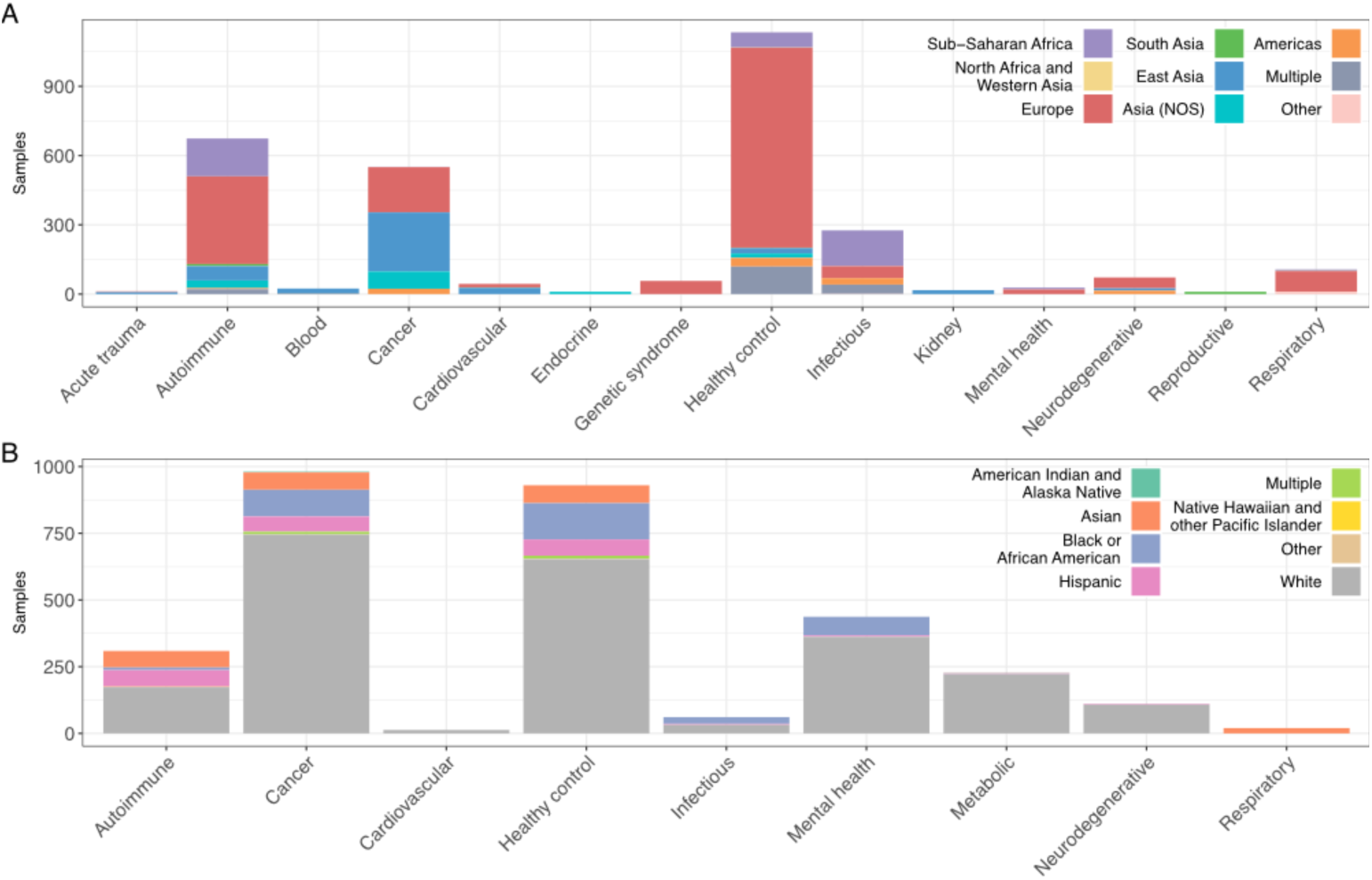
Number of individuals by population descriptor term and disease in SRA. Number of samples per disease or health condition (x-axis) broken down by A) Geographic/ancestral descriptors and B) US Census terms. Both panels focus on diseases with over 10 entries in SRA. A version of this plot showing all disease or health condition terms is available as **Figure S10**.

### Trends in reporting and bias over time and comparison to other data type

The number of RNA-seq samples in both dbGaP and SRA is rapidly increasing over time (**Figure S13**), including when focusing specifically on samples from underrepresented groups (**Figure 5**). However, the overall reporting of population descriptor terms has not improved with time. In SRA, year is not correlated with the proportion of samples that included any author-provided information on US Census terms, although this relationship trends in the expected direction (Pearson correlation: r = 0.45, p = 0.165), or with geographic/ancestral descriptors (Pearson correlation: r = −0.59, p = 0.075). In dbGaP, year does not significantly predict the proportion of samples with paired genotype data used to empirically derive population descriptors (r=0.28, p=0.35). The proportion of data derived from underrepresented groups in each public database has also increased over time, with year trending toward being positively correlated with the proportion of non-European ancestry individuals in SRA (r = 0.29, p = 0.106), although not with the proportion of non-white BioSamples in SRA (r = 0.09, p = 0.366) or non-European samples in dbGaP (r < 0.01, p = 0.996) (**Figure 5**). Thus, while the quantity and quality of reporting surrounding who is being included in public RNA-seq studies is still overall quite poor, attention to this issue does seem to potentially be improving over time in SRA.

**Figure 5.**
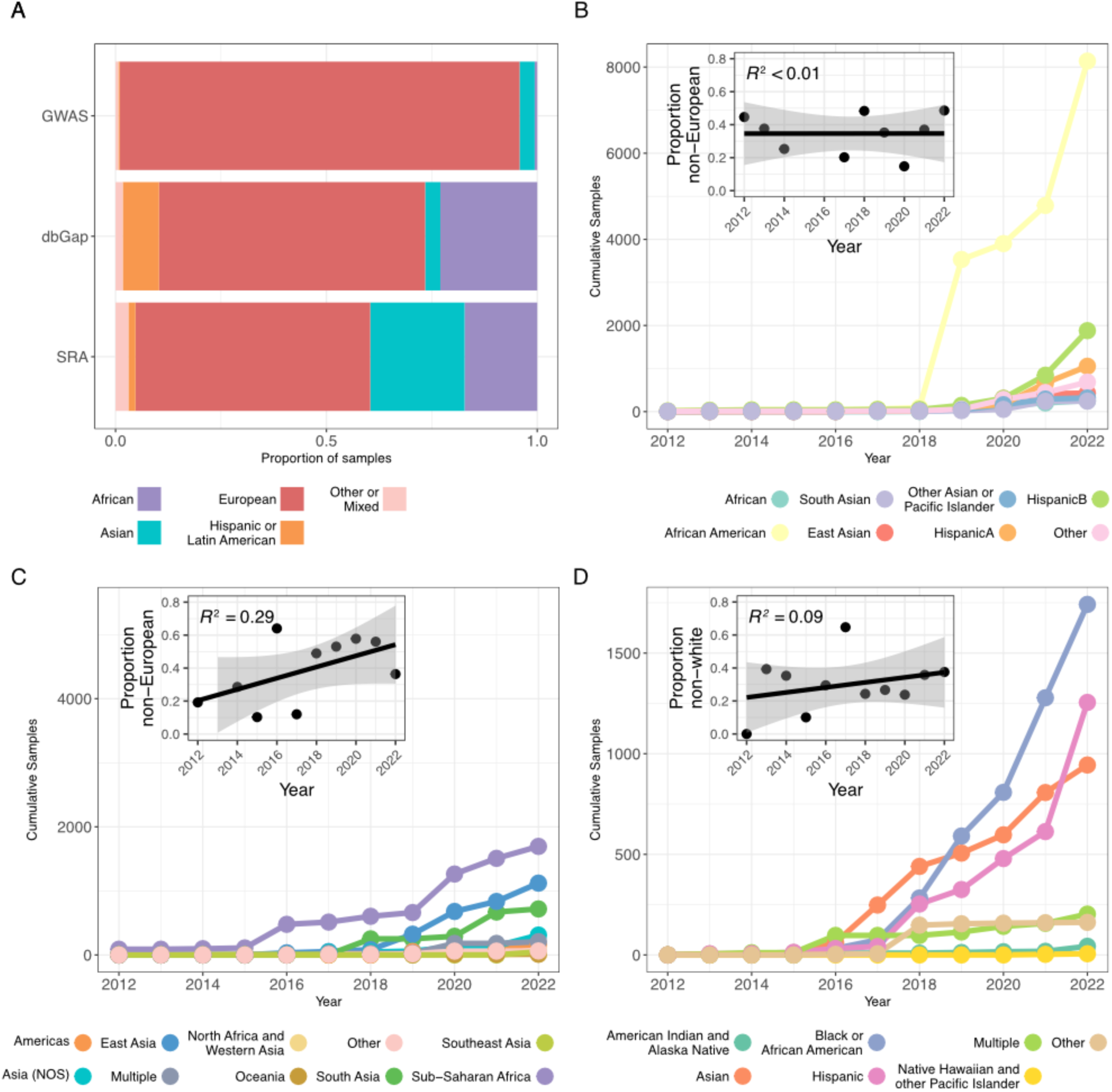
A) Proportion of samples by label in SRA, dbGaP and GWAS B) Cumulative samples over time by label in dbGaP (excluding European); inset shows the proportion of deposited samples each year not of European ancestry. C) Cumulative samples over time by geographic/ancestral label in SRA (excluding European); inset shows the proportion of deposited samples each year not labeled as European D) Cumulative samples over time by US Census label in SRA (excluding white); inset shows the proportion of deposited samples each year not labeled as white.

Relative to other public ‘omics data types for which similar underrepresentation estimates have been attempted, public RNA-seq data is in some ways more diverse but in other ways less diverse. For example, previous work estimated that 40% of public microbiome samples come from the US, where our estimates of bias for public RNA-seq samples are much higher (62-67%). We observe similar trends when grouping across high income countries more broadly, with 74% of microbiome samples coming from HICs in comparison to 94-100% of RNA-seq studies. In contrast, when we downloaded data from the GWAS diversity monitor, which breaks down individuals included in GWAS studies in terms of their continental ancestry, we found greater diversity of RNA-seq relative to GWAS participants (**Figure 5**). For example, 95% of GWAS participants are described as European, much more than the 56% and 63% estimated for SRA and dbGaP, respectively. Further, while year trends toward being positively correlated with the proportion of non European ancestry participants being added to public RNA-seq datasets, this does not appear to be the case for GWAS and the trend is instead in the opposite direction (r=-0.46, p=0.21; **Table S10**).

## DISCUSSION

Diverse sampling of individuals from globally representative groups is essential to robustly understand human genetics, evolution, and disease biology, and to combat health disparities. While some concerted, large-scale efforts are being made to characterize genetic diversity in undersampled groups (e.g., within Africa [22,23], Asia [24,25], and Central and South America [26]), less attention has been paid to biases that may exist in functional genomic datasets. Here, we analyzed diversity and bias in population descriptor terms within the two most popular public databases for RNA-seq studies, SRA and dbGaP. By analyzing RNA-seq data from multiple databases using multiple methods, we demonstrate that population descriptors are often incorrectly reported or not reported at all, and if they are, they are biased toward high income countries and individuals who are white or of European ancestry.

First, we found that very few studies report population descriptors for data either when uploading to public repositories, or in published papers. While there are valuable reasons for not reporting population descriptors [27], this trend was most pronounced in studies uploaded to SRA, where population descriptors are optional to report; as a result only 3% of individuals had any information provided (**Table 1**). The lack of information about population descriptors inhibits other researchers from using public data in informed ways, potentially biasing future secondary analyses in myriad ways [10]. Indeed, studies of LCLs from the 1000 Genomes Project have generally shown that genetic distance scales with the amount of differential expression detected between populations [13,28]; similarly, gene expression comparisons at the single cell level continue to identify inter-population differences that can be ascribed to both genetic variation, as well as interactions between genotype and environmental experiences that are stratified by population [25,29]. Although methods to infer genetic ancestry directly from transcriptomic data are becoming widespread [30,31], poor and incomplete reporting of population descriptors will make it challenging to design robust and unconfounded secondary analyses of public data. Such confounding has occurred in GWAS; for example, polygenic selection analyses on height that were overestimated due to uncorrected population stratification in the dataset [32–34]. Large meta-analyses of RNA-seq data could be similarly biased or underpowered if descriptors are not accounted for, and especially when used to disentangle genotype-by-environment interactions.

Second, we found that there is a heavy skew in RNA-seq data generation and deposition towards HICs, white individuals, and individuals of European ancestry. Analyses of microbiome, DNA methylation, and GWAS studies recapitulate these trends [3,5,6]. However, although RNA-seq studies have a higher bias of participants from the US when compared to microbiome studies, they have a higher diversity of ancestry groups than GWAS studies [3,6]. The dearth of diversity in all of these data types could limit the benefits of such studies. In RNA-seq studies in particular, associations between the transcriptome and disease have been found to vary across populations [14,35]; therefore, associations found among oversampled populations may not be valid in populations that are only sparsely sampled. Gene regulatory responses to infection, for example, show ancestry-associated differences between individuals with African versus European ancestry for ~10% of macrophage-expressed genes [14]. Furthermore, diseases that are studied in association with transcription are often inherently biased toward diseases present in the geographic region of the sampled individuals [36]. For example, relatively little is known about how tropical diseases impact gene expression, simply because they often are not present in HICs or among individuals who are white or have European ancestry.

Finally, we find that the reporting of population descriptors and inclusion of underrepresented groups in public RNA-seq databases has generally trended in the right direction over time, seemingly at a faster rate than non-European ancestry individuals are being included in GWAS. One potential explanation for this trend is that GWAS need much larger sample sizes relative to most analyses focusing on RNA-seq data [37]. Thus, improving representation in RNA-seq studies, which provide complementary information to what can be learned from genetic variation, may offer a more affordable route to bridging the biases and disparities in global datasets. We also see that certain tissues, such as blood samples, encompass more diverse participants relative to other types, probably because they are relatively easy to collect.

Our study has some limitations. The first is metadata quality: for the samples that did have population descriptors, race, ethnicity, and ancestry were often conflated (when following categories defined by the U.S. Office of Management and Budget for race and ethnicity, and geographic ancestry as described by [9]). Category and term usage also varied across countries, with 58.8% of samples deposited into SRA by institutions outside the US being labeled with geographic or ancestry-based descriptors (as reported by the data submitters) and 87.5% of samples deposited by US institutions labeled with racial descriptors (again, as reported by study submitters). Similar percentages were observed in dbGaP and the manual review. This reflects global differences in the usage of population descriptors—for example, “white” and “black” are racial descriptors in the US, but ethnic descriptors in the UK—and, likely, confusion amongst researchers as to the differences between descriptor categories (**Table S1**). Furthermore, in publications there is no standard way to report population descriptors and many papers bury this information in the supplement or present it in difficult to parse ways.

Consequently, we found that when multiple individuals manually reviewed the same SRA accession, there were inter-individual differences, highlighting the difficulty in identifying key information (**Table S11**). A final limitation is that we only conducted analyses in SRA and dbGaP, and it is possible that other databases have different trends of over- and underrepresentation. For example, it is important to note that while dbGaP data is behind access controls, SRA data is freely available, making it challenging to maintain data sovereignty, guarantee the genetic privacy of donors, or to ensure any future use of the data is consistent with the initial informed consent conditions; these conditions can make researchers working with underserved and underrepresented communities less likely to deposit data there [38,39].

Practically, our study illustrates several ways in which the reporting of population descriptors and general diversity of RNA-seq data in publicly available datasets is lacking. Stakeholders of resource databases should consider how to incentivize standardized reporting. Other studies have addressed barriers that impede progress to increased diversity in ‘omics studies, such as lack of diversity in researchers themselves, funding [10], and legacies of harm and distrust [7,8]; a major ongoing conversation in human genomics is how to improve global representation in our studies, including RNA-seq studies specifically [1,10]. Doing so is mission critical for all fields, because the current focus on white or European ancestry individuals living in urban settings in HICs provides only a narrow representation of both human biology and the range of social, cultural, and physical environments occupied by our species. This significantly limits our collective ability to understand human evolution, diversity, and environmental responses, as well as to move beyond initial discovery to equitable clinical outcomes.

## METHODS

### Generating a list of all human RNA-seq SRA projects

We generated a list of all publically available human RNA-seq projects in SRA using the recount3 database [40]. Specifically, we used the recount3 Study Explorer (https://jhubiostatistics.shinyapps.io/recount3-study-explorer/) to generate a list of all public human RNA-seq SRA projects with at least 10 BioSamples (n=8,742). Because recount3 was published in 2021 and therefore does not include studies from more recent years, we also used search queries provided by the recount3 authors to gather projects from more recent years. Specifically, we used the SRA Run Selector Tool and the following search parameters executed on 14-Sep-23:

((((((((illumina[Platform]) AND rna seq[Strategy]) AND transcriptomic[Source]) AND public[Access]) NOT size fractionation[Selection]) AND human[Organism])) AND (“2019/01/01”[Publication Date] : “2022/12/31”[Publication Date])).

We merged these results with the recount3 dataset, filtered for unique studies and studies with at least 10 BioSamples, and arrived at a final list of human RNA-seq SRA projects that spanned 2012 through September of 2023. Because the 2023 data do not represent an entire year, for analyses of trends over time, we excluded 2023.

### Extracting and processing metadata for SRA projects

NCBI hosts the SRA BioSample database which can be accessed via the SRA Run Selector online tool. This database includes information about all raw sequencing data deposited into SRA [41]. BioSample metadata is deposited by researchers at the time of upload to the SRA public repository. There are nine mandatory columns that must be completed by all submitting studies, including “Center Name” and either “Cell line” or “Cell type”, but SRA encourages individual researchers to create additional columns to fully capture the structure of their data. Reporting of additional descriptors is at the discretion of the authors and can be written in or derived from a list of 485 possibilities (https://www.ncbi.nlm.nih.gov/biosample/docs/attributes/). These additional descriptors are not curated for consistency with already deposited data. Thus, the “Population” column contains, for example, population descriptor terms such as “YRI” or “GBR”, referring to the Yoruba and British from England and Scotland human populations sampled by the 1000 Genomes Project [42], but also FACS gating results such as “POU3F2 High” or “Vd1 CD27HI”, representing sorted cell populations.

We downloaded all available metadata for the human RNA-seq projects included in our list (derived from recount3 as well as our own parallel queries, as described above). In total, SRA metadata for these projects included 1,557 columns, the vast majority of which were only used by a small subset of studies. From these, we manually identified columns containing information that would help us understand sources of study design features and biases. We identified eight columns consistently used for reporting of population descriptor terms across studies (“ancestry”, “DONOR_ETHNICITY”, “ETHNICITY”, “Population”, “primary_race”, “RACE”, “race.ethnicity” and “reported_race”). We also identified six columns (“tissue”, “tissue_type”, “Organism_part”, “cell_type”, “source_name” and “Cell_line”) that contained information on the assayed cell or tissue type, and five columns that contained information on any disease or health condition associated with the samples (“disease”, “disease_state”, “Diagnosis”, “health_state” and “DONOR_HEALTH_STATUS”). Finally, we used the mandatory Center Name column, which contains the name of the institution submitting data to SRA, to infer the likely country of the study team, although we note this may not necessarily reflect where samples were collected.

Data can be submitted to SRA directly or indirectly, via the Gene Expression Omnibus (GEO); in the second case the Center Name is reported as “GEO” in SRA. For those cases (123 studies) we used NCBI’s Entrez Programming Utilities (E-utilities), as implemented in the package rentrez [43], to identify the GEO BioProject associated with the SRA Study ID and retrieved the Center Name in this way. We used a similar approach to identify the depositing institution for all dbGaP studies.

### Harmonizing terminology in SRA project metadata

Once we had identified relevant columns for all metadata, we proceeded to harmonize and backcode the descriptors in each dataset. First, all data was manually inspected to control for misspellings, acronym use, and the use of incorrect/uninterpretable terms (e.g., “Race: Human”). Cell type and tissue information was further standardized and grouped by searching terms against the Cell (CL) and UBERON ontologies [44,45]; disease information was standardized and grouped using the Disease Ontology [46]; in all cases we resolved any ambiguities by manually inspecting the SRA Study summary, and if data could not be disambiguated that way, we coded the relevant column as “NA”. Center Name was mapped to country via manual internet searches, and countries were grouped using World Bank country and lending groups data (downloaded on 23-April-2024 from: https://datahelpdesk.worldbank.org/knowledgebase/articles/906519-world-bank-country-andlending-groups).

Given ambiguities in term usage, harmonization and back-coding of population descriptors was more challenging. First we identified two different types of widely-used descriptors: geographic/continental ancestry descriptors and US Census racial/ethnic descriptors. After manual inspection, we considered any entries in the “ancestry”, “ETHNICITY” and “Population” columns to represent a geographic/ancestry descriptor, and any entries in “DONOR_ETHNICITY”, “primary_race”, “RACE”, “race.ethnicity” and “reported_race” to be primarily US Census racial/ethnic descriptors. We then harmonized terms in either category using the US Census categories for race and ethnicity (American Indian or Alaska Native, Asian, Black or African American, Native Hawaiian or Other Pacific Islander, White, and Hispanic) and geographic groupings (Sub Saharan Africa, Middle East and Western Asia, South Asia, Southeast Asia, East Asia, Asia (not otherwise specified), Europe, Oceania, Americas); these categories match previously published studies, but provide additional granularity in Asia, which is home to over half of the world’s population. Importantly, we deferred to author-supplied metadata categories for grouping sample terms even when the reported terms were ambiguous or appeared to be incorrectly categorized. For example, a sample labeled as “Ethnicity: Black or African American” was coded as “Geography: Subsaharan Africa” while “Race: Malay” was changed to “Race: Asian”.

Following this, 9,825 samples were associated with only a geographic term, 13,668 samples were associated with a race term, and 535 samples were associated with both geographic and racial terms. This dual coding of a subset of samples was due to either submitters duplicating information in the “RACE” and “ETHNICITY” columns (e.g. SRP068551 reports ethnicity and race both as “White or Caucasian”) or providing complementary information in both columns (e.g., SRP292867 reports ethnicity as “Korean” or “German” and race as “Asian” or “White”, respectively). To avoid double-counting samples, and because the pertinent SRA columns primarily reflected US-typical usage of US Census racial terms, we retained the racial term if the Center Name was that of an US-based institution, and the geographic term otherwise.

The only exception to this overall approach was in identifying Hispanic samples. This is the only ethnicity recognised by the US Census, and is nested one level above all racial terms (i.e., Hispanic individuals cannot report a race, and only non Hispanic individuals can report a race; see https://data.census.gov/table?q=United+States+Race+and+Ethnicity). Because of this, Hispanic status was frequently reported in either the “ETHNICITY” and “RACE” columns, and we therefore considered a sample to be Hispanic regardless of the metadata column this information was found in. Furthermore, because current US Census practice is that the Hispanic label supersedes racial categories, we coded all samples labeled as Hispanic that also reported racial information as exclusively Hispanic. This led to 13 studies containing a mixture of samples with either a geographic or a US Census label, but we chose not to alter this to avoid further manual reclassification.

We also considered an alternative approach in which we treated terms that unambiguously matched US Census categories as racial descriptors regardless of metadata category (for example, a sample labeled as “Population: Native American and Alaska Native” was recoded to “Race: Native American and Alaska Native”) and likewise reclassified some racial terms as geographic when additional metadata suggested the original usage was not aligned to that of the US Census: for example, both Brazil and Singapore use race as a means of classifying individuals, but given the culture-specific nature of race, these should not be mapped to US terms even if the term is the same, e.g. “White”. We found that this alternate approach required more arbitrary decision making, relied on our pre-existing knowledge of how different countries describe individuals, and had little significant impact on our general conclusions.

The full lists of terms associated with each metadata descriptor, and harmonized groupings used in all analyses, are available as **Table S5**.

### Extracting and processing metadata from dbGaP

We downloaded the metadata from all 649 RNA-seq studies (167,389 samples) stored in dbGaP as of April 26, 2024 (by selecting “RNA-seq” and “RNA Seq (NGS)” studies in the dbGaP advanced search portal). Studies in dbGaP that include genotype data (28% of the original 167,389 samples) are automatically run through the GrafPop program pipeline (Genetic Relationship And Fingerprinting) and assigned to continental ancestry groups [21,47]. More specifically, GrafPop calculates genetic distances from each sample relative to several reference populations and estimates ancestry groupings based on cutoffs/discrete groups of these distance metrics (https://www.ncbi.nlm.nih.gov/projects/gap/cgibin/GRAF_README.html). The terms used in the dbGaP metadata for potential ancestry groupings are: Other, Hispanic1 or 2, Other Asian or Pacific Islander, European, African American, South Asian, East Asian, and African. Hispanic 1 and 2 correspond to different arbitrary cutoff points for groupings as described in the GrafPop documentation. We note that, in the documentation, Hispanic 1 and 2 (ethnicity terms) are replaced with Latin American 1 and 2 (ancestry terms). Further information about GrafPop groupings are provided in **Table S2**.

### Manual review of publications associated with SRA depositions

RNA-seq SRA accession numbers were randomized and reviewed for manual data extraction by the study authors. We reviewed 620 total SRA accession numbers. 58 of the 620 SRA accession numbers were reviewed a second time to understand inter-coder consistency (see next section). Of the repeats, one entry was randomly chosen to include in the analyses presented in the main text to avoid double counting any studies.

For each randomly chosen SRA accession for manual review, the study was excluded from further analyses if it met any of the following criteria: (i) studies only used established, cancerous, or cancer-derived cell lines, (ii) studies that did not measure gene expression, (iii) studies using only non-human samples, (iv) studies involving genetic manipulation, or (v) the project lacked an associated paper describing the methods and data generation. After screening, a total of 321 studies met our criteria for further analysis. For the 312 studies, we then extracted information about individual study participants by sample type (i.e., primary cells, LCLs, fibroblasts, or iPSCs). Specifically, among other information, we identified (i) the number of individuals, (ii) any population descriptors used, (iii) the number of individuals reported for each population descriptor, (iv) sampling location (country), (v) country of last author institution, and (vi) if genotypes were generated. These are the pieces of information that were most heavily used for downstream analysis, but **Table S3** describes all metadata extracted during manual review. A total of 16,801 individuals were included in the 312 studies that we evaluated.

Population descriptors were recorded as reported by the study author to maintain consistency (e.g., “Race as reported by the study authors”, “Ethnicity as reported by the study authors”). In cases where the population descriptor term was not explicitly defined by the authors, we attempted to infer it based on the labels used. To standardize the sample-specific labels for each study, we used the same process described for the SRA metadata backcoding to assign geographic/continental ancestry or US census race/ethnic categories.Thus, studies with the following responses were considered to fall under geographic groupings: “Ancestry as inferred by me from the study” and “Ancestry as reported by the study authors”. Similarly, studies with the following responses were considered to fall under US Census racial/ethnic categories: “Ethnicity as inferred by me from the study”, “Ethnicity as reported by the study authors”, “Race as inferred by me from the study”, and “Race as reported by the study authors”.

During the manual review of the studies, the country that the samples were collected in was reported for 2,947 individuals (18% of the total). For downstream analyses, we used this country information to determine income group (low, lower-middle, upper-middle, and high) as defined by the World Bank (**Table S6**).

### Quantifying the difficulty of parsing population descriptors during manual review

A single individual repeated the manual review for 51 SRA accessions to understand repeatability (**Table S11**). Of the 51 repeated reviews, 30 agreed on including the study in the manual review based on the exclusion/inclusion criteria described above. We used this subset to further review consistency of reporting.

We found that reviewers mostly agreed on sample types (“Primary samples”, “LCL, fibroblast or iPSC cell lines”) and identifier labels. For example, based on the answer to “Sample type”, we found that there were 25 entries that agreed the study included “Primary samples”.

However, there were more disagreements when recording the number of individuals, if/what type of population descriptors were reported (“Ancestry”, “Ethnicity”, “Race”, “not reported”), and sampling location. In particular, when looking for the number of individuals, discrepancies may have arisen if the paper reported numbers or population descriptors for an initial cohort, but were not clear when a subset was used for RNA-seq; furthermore, samples may have been discarded during the methods due to poor quality or other filtering reasons, often leading to small discrepancies in the number of samples reported by the two independent reviewers.

Information about samples and individuals were also recorded by authors in diverse ways: they were often reported in a variety of locations within the paper, including the methods, results, figures, tables, supplementary information documents or externally linked datasets (and these pieces of information were not always internally consistent). Overall, as there was no standard way of reporting this information, this could lead to ambiguity and thus inter-individual differences in how information is interpreted. However, we note that coding differences between individuals was relatively minor (e.g., numbers were generally within the same order of magnitude or representative of the same biases in the study), even if individual coders sometimes disagreed about which version of sample sizes/specifics to extract from a given paper.

### Analyses of over- and under-representation

In general, analyses of over- and under-representation compared country- or term-specific proportions observed in the metadata to proportions obtained from the US Census (**Table S4**) or the United Nations (**Table S7)**. For the overrepresentation data presented in **Figure S3**, we first calculated the number of individuals living in each region using country population data from the UN (**Table S7**). Since large numbers of individuals from European and sub-Saharan African ancestries live outside these regions, we adjusted these totals by taking into account the number of individuals living overseas from either group (as reported in the Wikipedia “European Diaspora” and “African Diaspora” pages, accessed on September 2, 2024) and adjusting all other regional population estimates downwards accordingly [48,49].

## Data and code availability

All raw data has been deposited in Figshare and is available at https://figshare.unimelb.edu.au/projects/RNA-seq_diversity/221698. Summarized versions of the data are available in the Supplementary Tables. All code used to analyze the data and generate figures is available at https://github.com/AmandaJLea/RNAseq_diversity.

## Supporting information

All supplementary tables

All supplementary figures

## ACKNOWLEDGMENTS

We thank all members of the Lea and Gallego Romero labs for the feedback and support, in particular Axel Cortada-Mccorkell and Lu McNaughton. AJL acknowledges research support from the Canadian Institute for Advanced Research (Azrieli Global Scholars Program), the Kinship Foundation (Searle Scholars Program), and the Pew Charitable Trusts (Pew Biomedical Scholars Program). IGR acknowledges research support from Australian Research Council Discovery Project DP200101552 and National Health and Medical Research Ideas Grant GNT2020501, and was partially supported by the European Union through Horizon 2020 Research and Innovation Program under Grant No. 810645 and the European Union through the European Regional Development Fund Project No. MOBEC008. St Vincent’s Institute acknowledges the infrastructure support it receives from the National Health and Medical Research Council Independent Research Institutes Infrastructure Support Program and from the Victorian Government through its Operational Infrastructure Support Program. AMA acknowledges research support from the National Science Foundation Graduate Research Fellowship Program under Grant No. 1937963 & 2444112.

## SUPPLEMENTARY INFORMATION

SI Tables: RNA-seq diversity - SI tables v2

SI Figures: RNA-seq diversity Supplementary Figures

