## Supplementary material for "Public RNA-seq data are not representative of global human diversity": All supplementary figures

**Figure S1: Under and over-representation of samples deposited from a given country relative to the estimated global population across all three databases.** X-axis represents the relative ratio of the proportion of samples deposited from a given country versus the proportion of people globally that live in that country according to the United Nations (see **Table S7**).

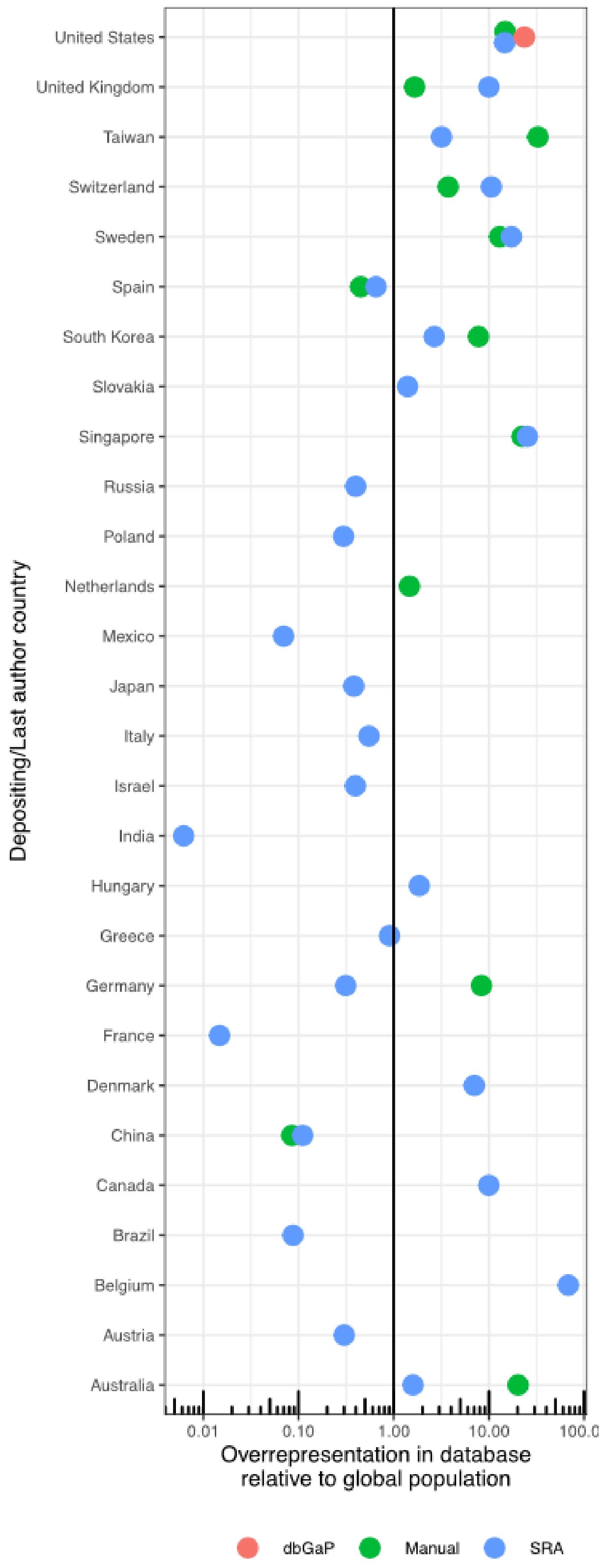

**Figure S2: Number of samples per SRA study broken down by World Bank Economic Region of the depositing institution.** Boxplots highlight the median, 25th percentile, and 75th percentiles. Regions are further detailed in **Table S6**.

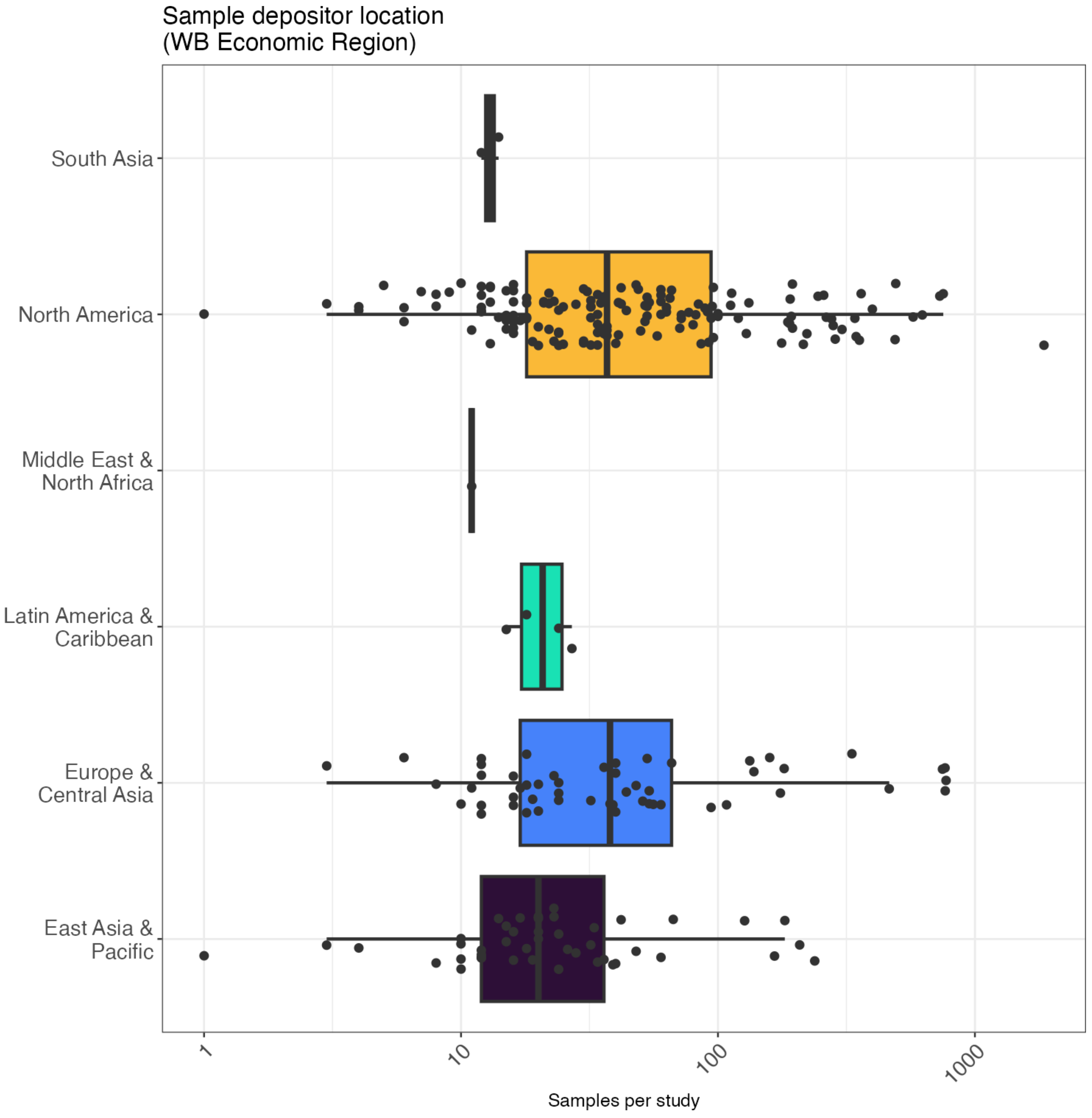

**Figure S3: Under and over-representation of samples by geographic/ancestral descriptor term.** X-axis represents the relative ratio of the proportion of samples from a given region versus the proportion of people globally that live in that region according to the United Nations (see **Table S7**).

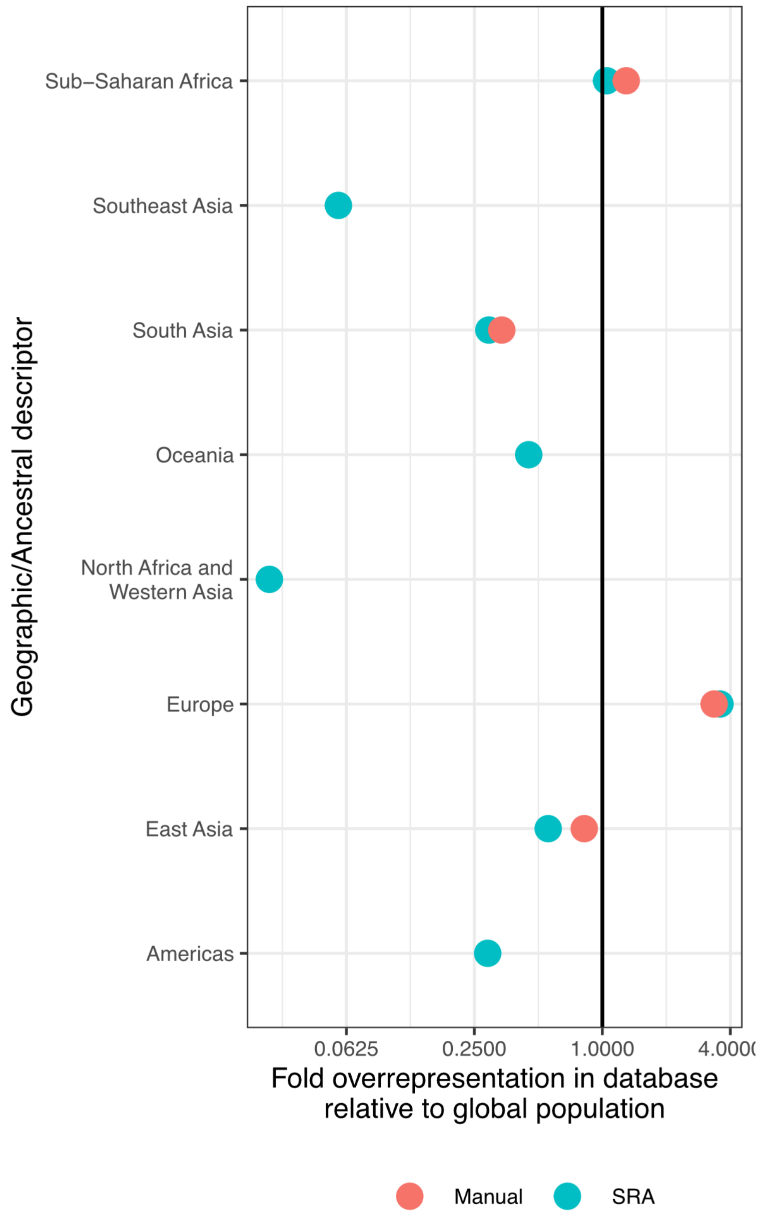

**Figure S4: Under and over-representation of samples by US Census term.** X-axis represents the relative ratio of the proportion of samples labelled with a given term versus the proportion of people in the US that identify with that term according to the US Census (see **Table S4**).

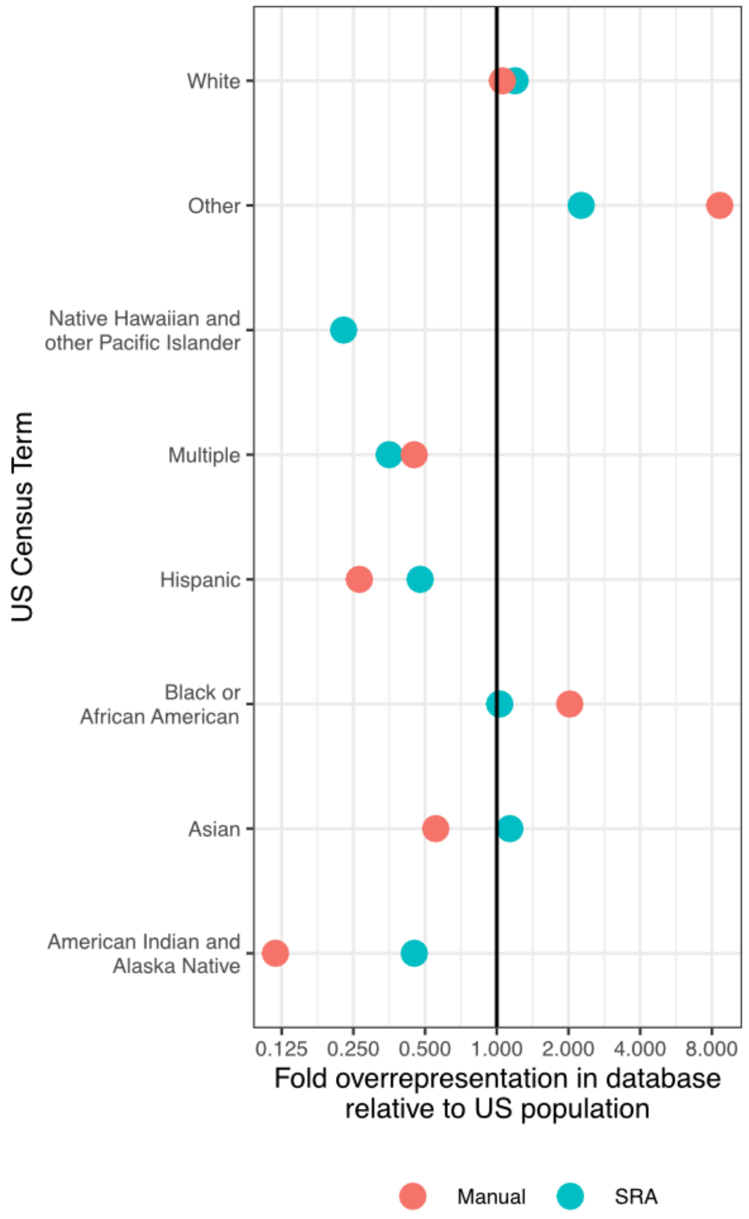

**Figure S5: Number of samples per SRA study broken down by population descriptor terms.** Boxplots highlight the median, 25th percentile, and 75th percentiles.

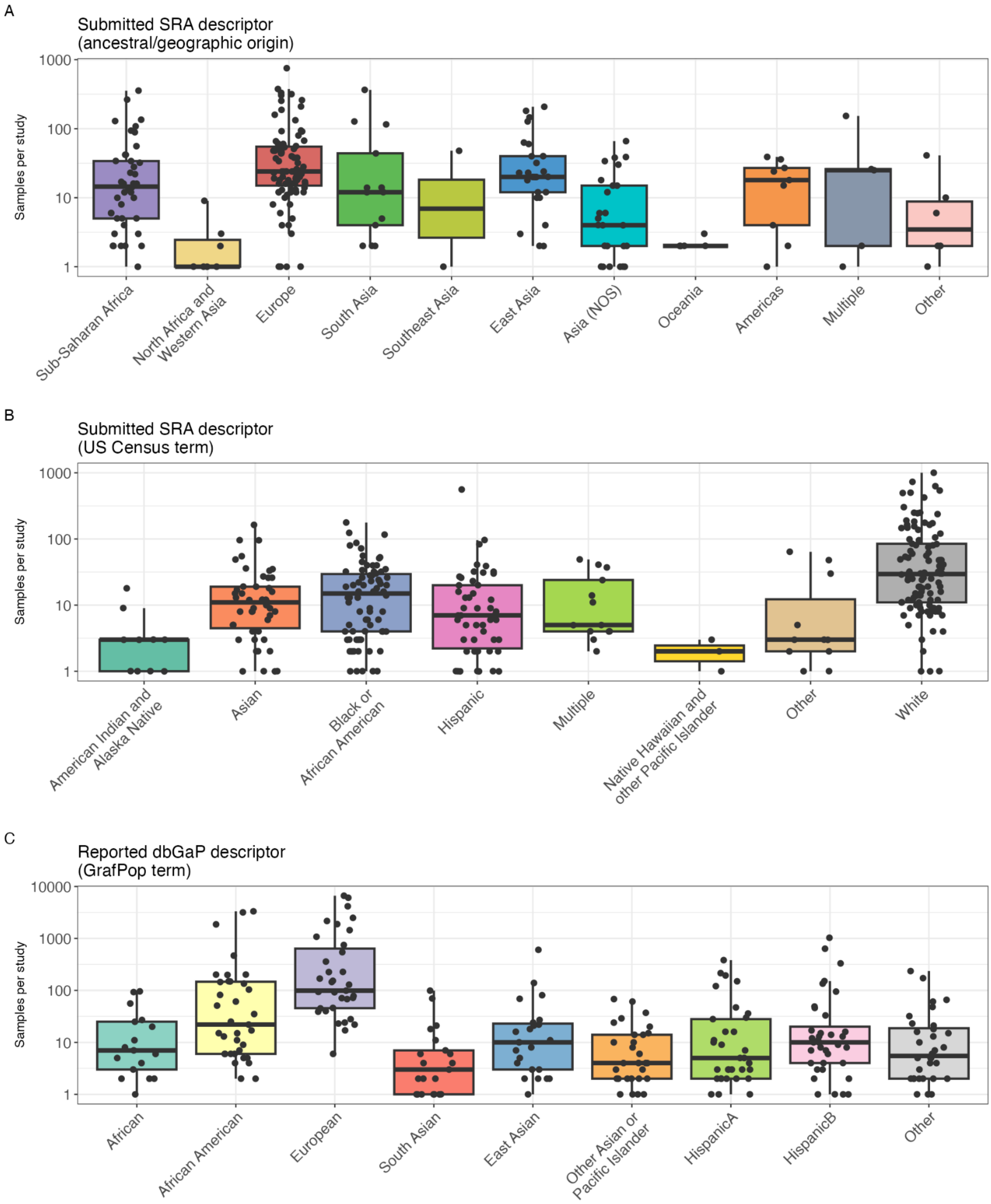

**Figure S6: Number of BioSamples in SRA broken down by population descriptor term and World Bank Economic Region of the depositing institution.** Breakdown is shown for A) geographic/ancestral descriptors and B) US Census terms.

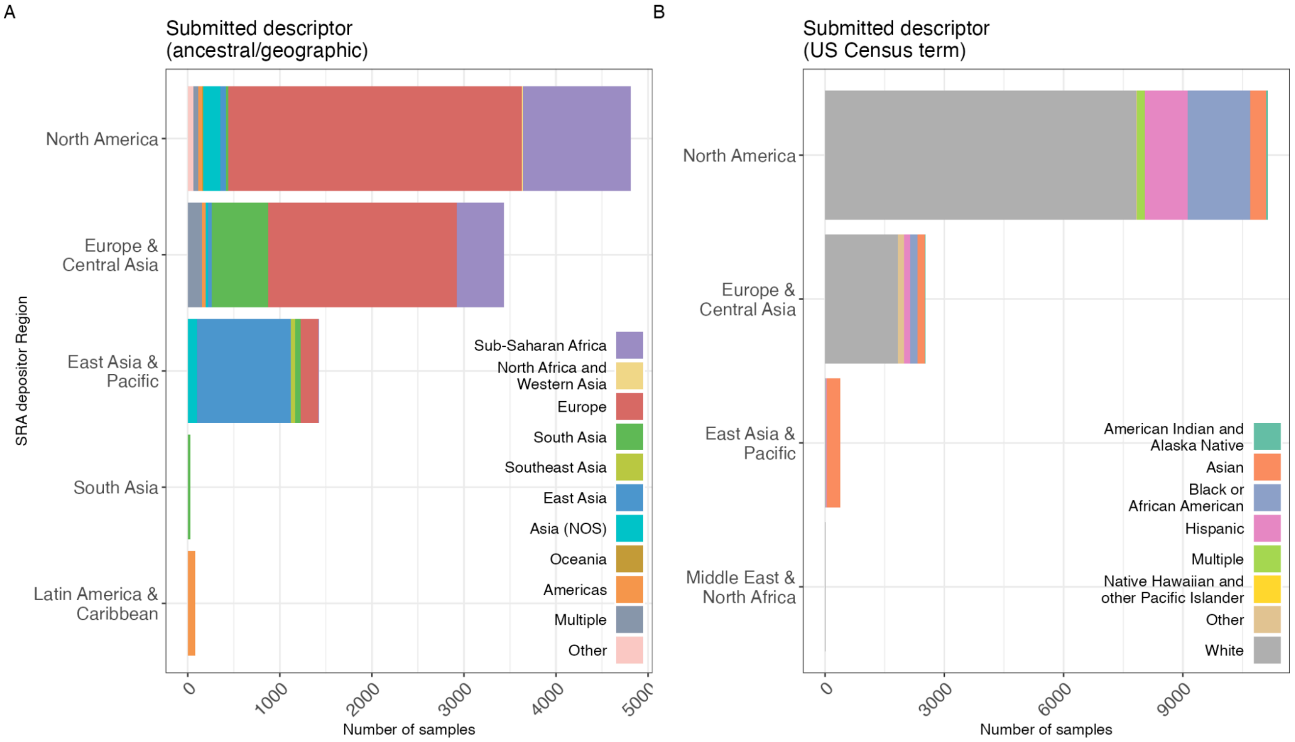

**Figure S7: Number of BioSamples in SRA broken down by population descriptor term and tissue.** Breakdown is shown for A) geographic/ancestral descriptors and B) US Census terms.

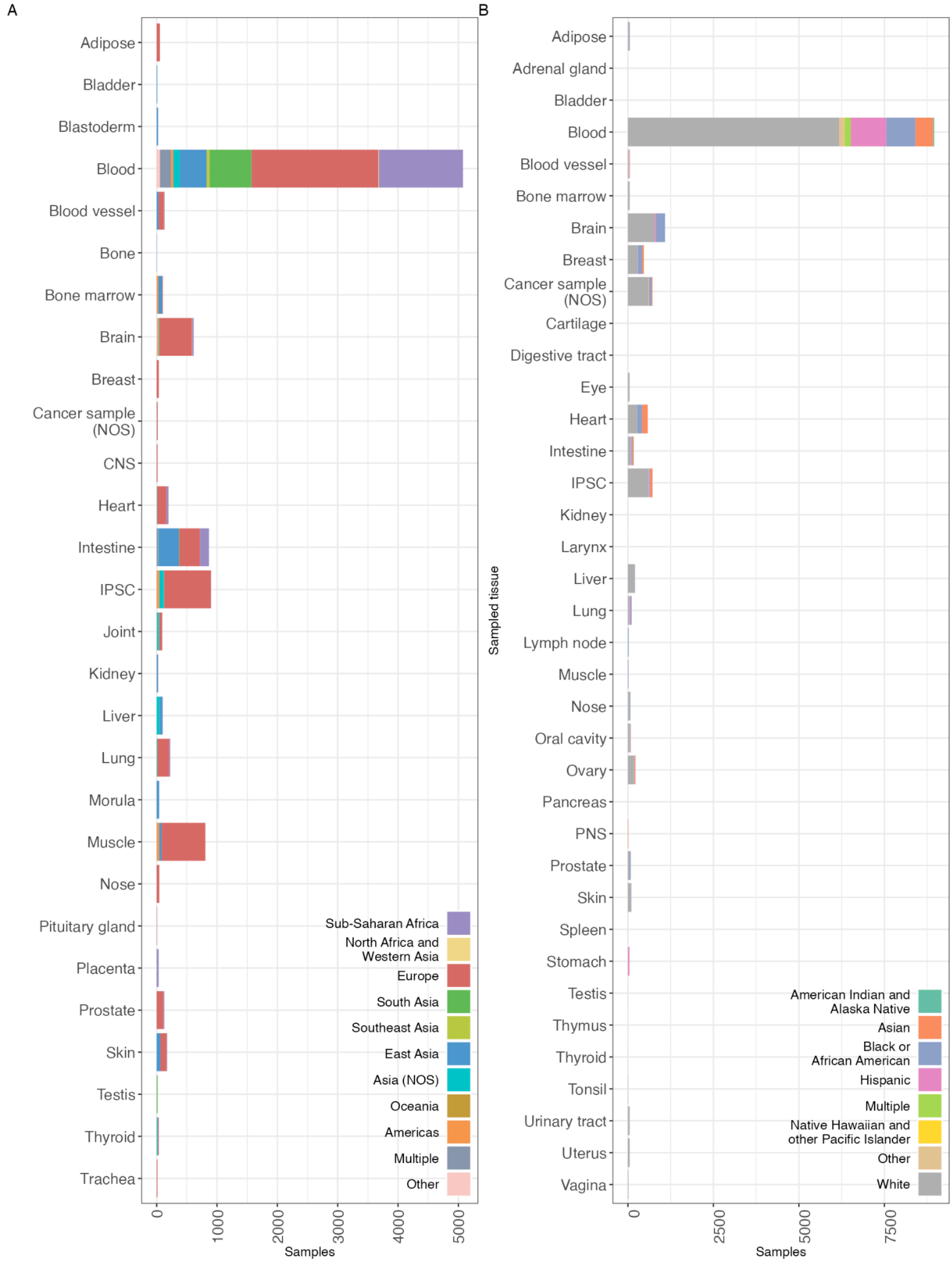

**Figure S8: Number of BioSamples in SRA broken down by population descriptor term, tissue, and World Bank Economic Region of the depositing institution** (as in Table S6).

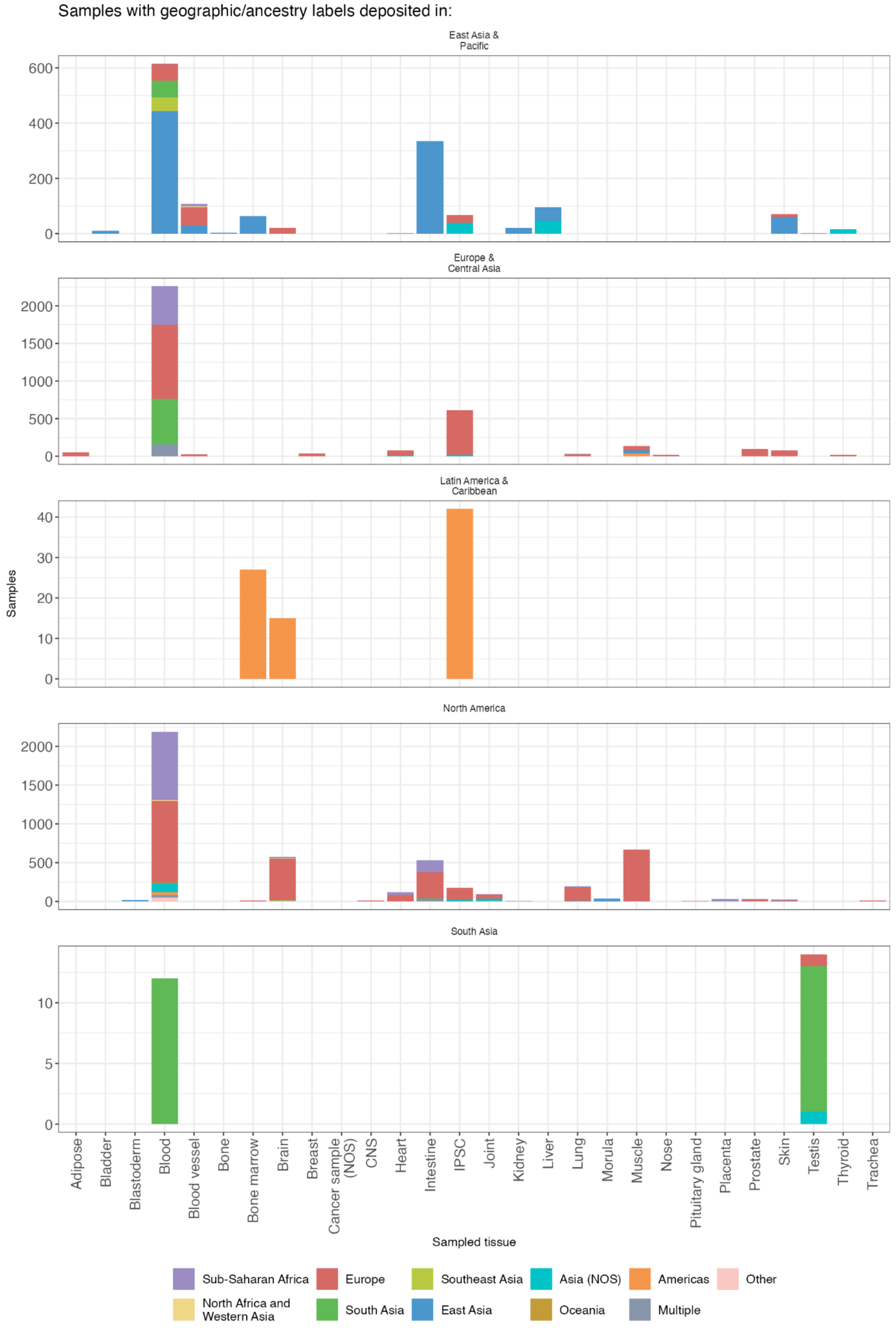

**Figure S9: Number of BioSamples in SRA broken down by population descriptor term, tissue, and World Bank Economic Region of the depositing institution** (as in Table S6).

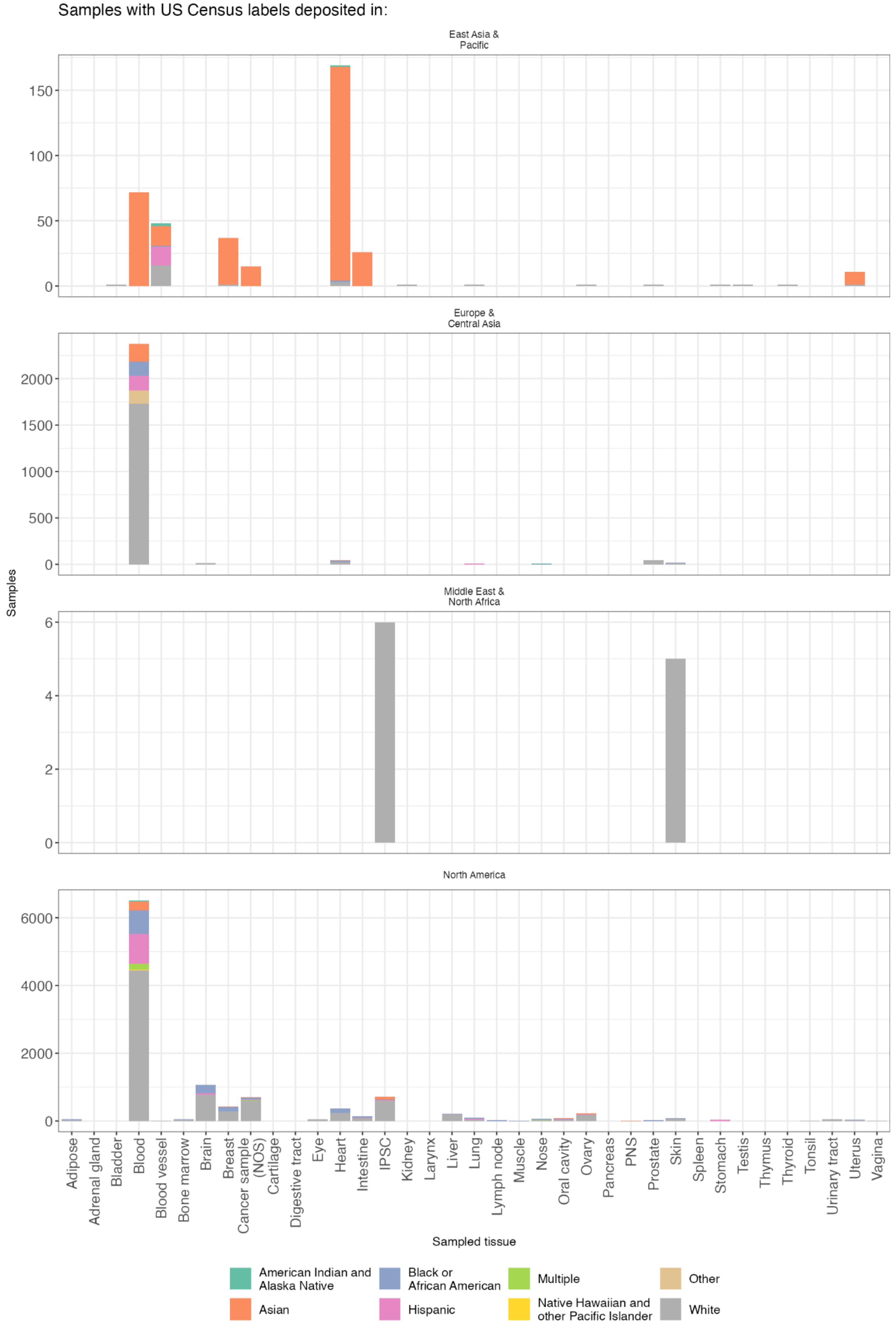

**Figure S10: Number of BioSamples in SRA broken down by population descriptor term and disease term.** Breakdown is shown for A) geographic/ancestral descriptors and B) US Census terms.

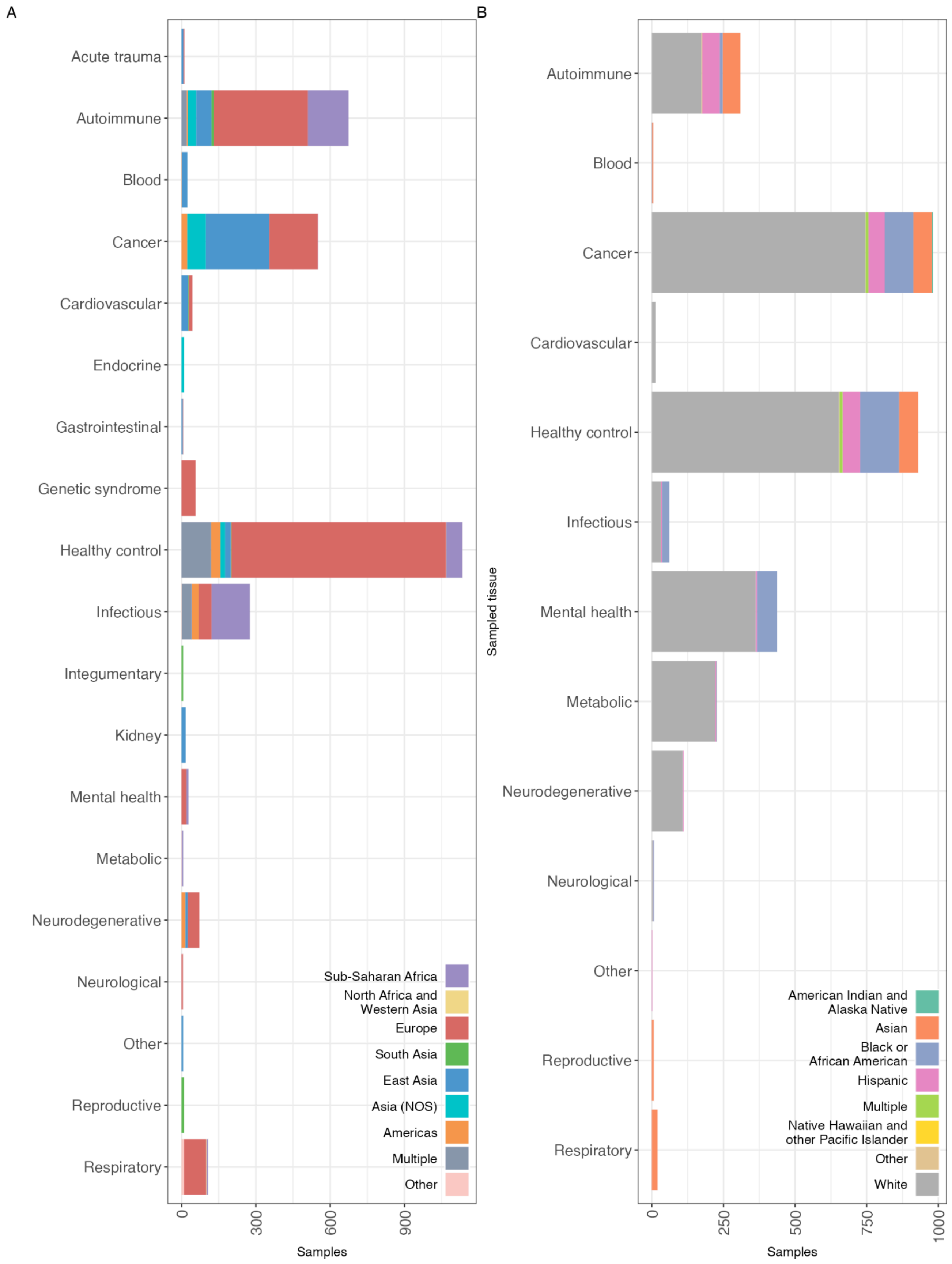

**Figure S11: Alluvial plot illustrating the relationship between disease focus, tissue of origin, and World Bank Economic Region of depositing institution for samples with disease focus information and geographic/ancestry descriptors in SRA. Each coloured flow represents connections between categories, with the width of the flows indicating the number of samples across categories. Flows are coloured by geographic/ancestry descriptors.**

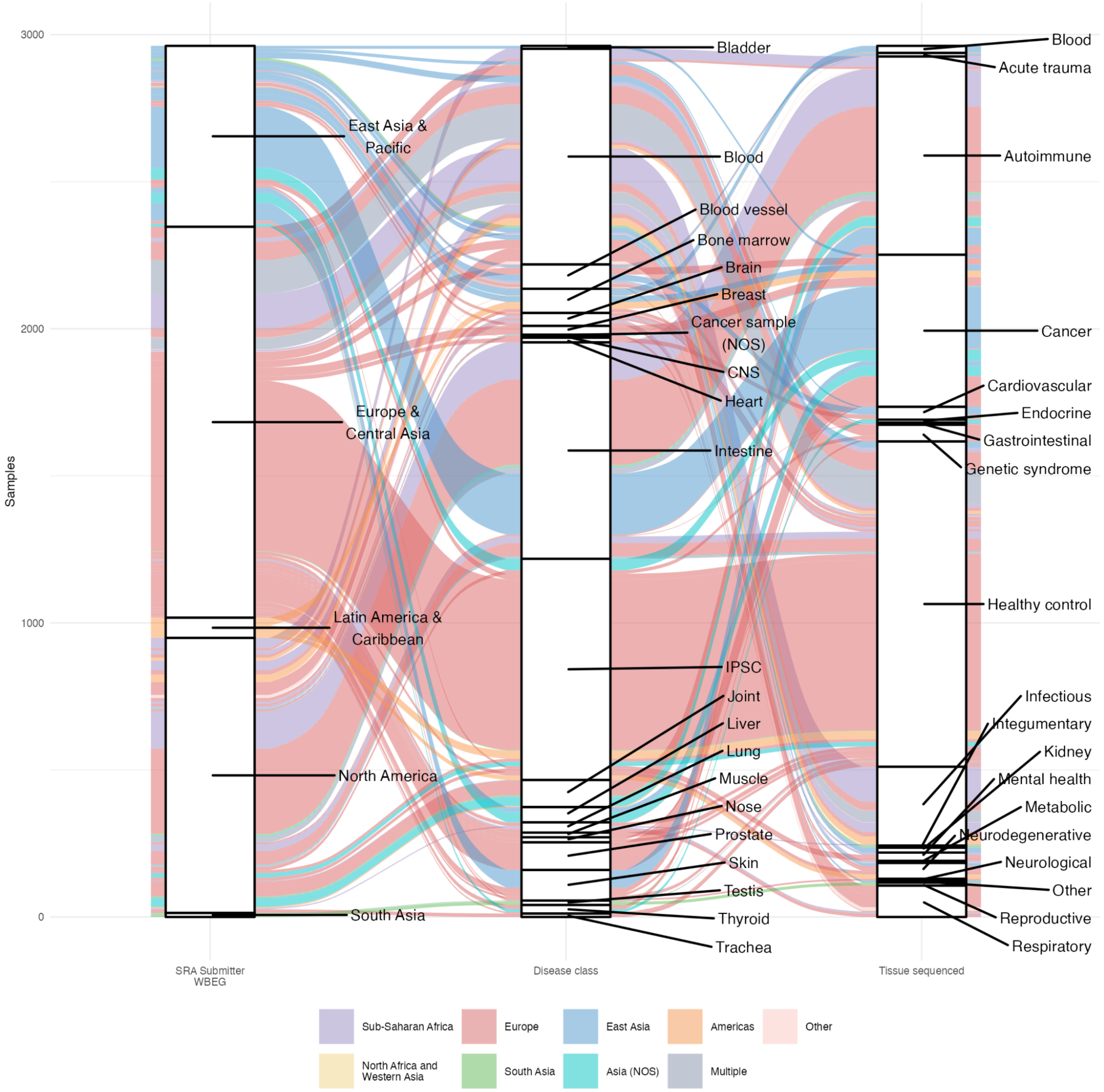

**Figure S12: Alluvial plot illustrating the relationship between disease focus, tissue of origin, and World Bank Economic Region of depositing institution for samples with disease focus information and US Census descriptors in SRA. Each coloured flow represents connections between categories, with the width of the flows indicating the number of samples across categories. Flows are coloured by US Census descriptor.**

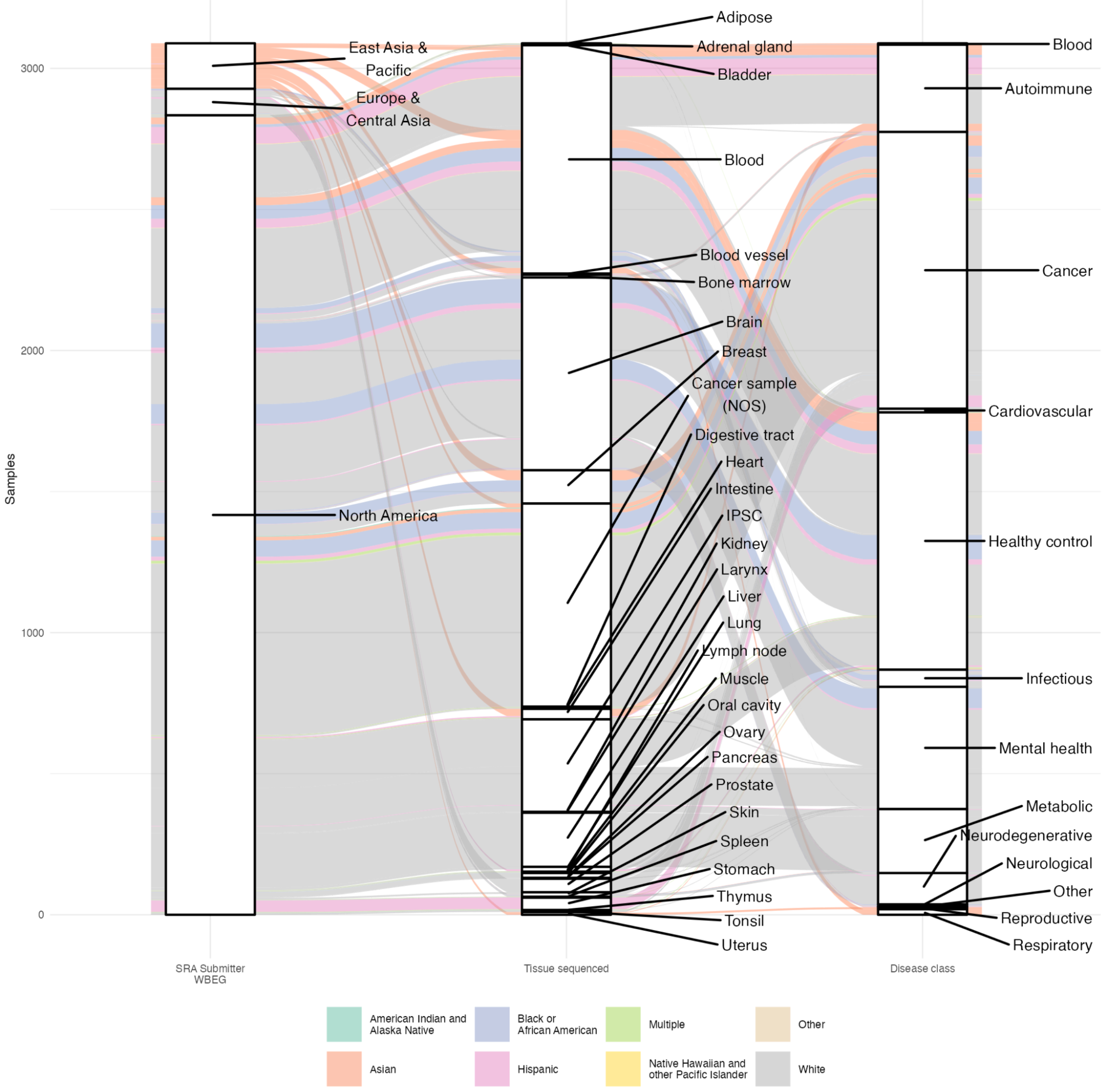

### Figure S13: Cumulative samples over time deposited to SRA and dbGaP

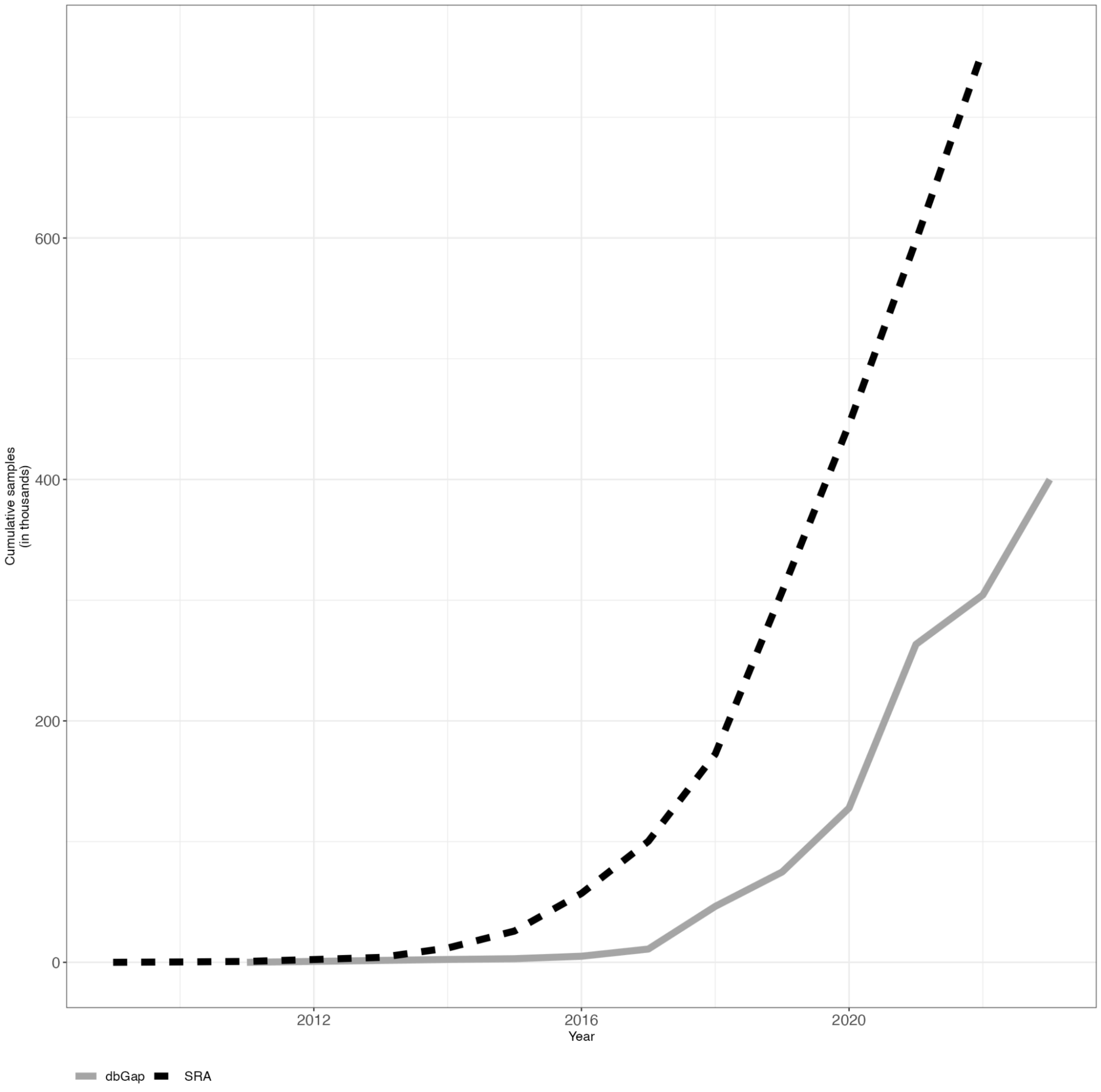
